# Ghrelin delays premature aging in Hutchinson-Gilford progeria syndrome

**DOI:** 10.1101/2023.05.02.539084

**Authors:** Marisa Ferreira-Marques, André Carvalho, Ana Catarina Franco, Ana Leal, Mariana Botelho, Sara Carmo-Silva, Rodolfo Águas, Luísa Cortes, Vasco Lucas, Ana Carolina Real, Carlos López-Otín, Xavier Nissan, Luís Pereira de Almeida, Cláudia Cavadas, Célia A. Aveleira

## Abstract

Hutchinson-Gilford progeria syndrome (HGPS) is a rare and fatal genetic condition arising from a single nucleotide alteration in the *LMNA* gene, which leads to the production of a defective lamin A protein known as progerin. The buildup of progerin hastens the onset of premature and expedited aging. Patients with HGPS exhibit short stature, low body weight, lipodystrophy, metabolic dysfunction, and skin and musculoskeletal abnormalities and, in most cases, die of cardiovascular disease by their early teenage years. Currently, no effective cure or treatment for the disease highlights the importance of discovering new therapeutic strategies. Herein, we present evidence that the hormone ghrelin, besides promoting autophagy and progerin clearance, rescued several cellular hallmarks of premature aging of human HGPS fibroblasts. Using an HGPS mouse model, *Lmna*^G609G/G609G^ mice, we also show that ghrelin administration rescued the short-lived mice molecular and histopathological progeroid features, prevented progressive weight loss at later stages, reverted the lipodystrophic phenotype, and extended lifespan. Thus, we disclose that modulation of ghrelin signaling may give rise to new treatment targets and translational approaches that may improve outcomes and the health quality of HGPS patients and natural aging pathologies.

## Introduction

Hutchinson–Gilford progeria syndrome (HGPS; OMIM#176670) is a rare and fatal genetic condition characterized by premature and accelerated aging, with an incidence of approximately 1 in 4–8 million live births (1, 2). Children with HGPS exhibit low body weight, lipodystrophy, abnormalities in the skin, musculoskeletal, and cardiovascular systems, and typically die from myocardial infarction or stroke at a median age of 14.6 years (3, 4). In the classical form of HGPS, a specific mutation (c.1824C>T,p.G608G) causes a single C-to-T transition point mutation at position 1824 of the *LMNA* gene. This mutation leads to the activation of a previously hidden splice site, resulting in the loss of the terminal 150 nucleotides of exon 11 (5, 6). The consequence of this deletion in the C-terminal domain is the formation of a truncated form of prelamin A, named progerin. Progerin does not undergo the normal post-translational processing retaining a toxic farnesyl modification. Permanent farnesylation causes progerin accumulation in the inner nuclear membrane, which is at least partly responsible for the HGPS phenotype (7-9).

Ghrelin is a 28-aa Ser3 acylated peptide, and its functionally relevant endogenous receptor is the growth hormone secretagogue receptor (GHS-R1a) (10). Both ghrelin and its receptor have a ubiquitous expression throughout the body (11, 12), supporting ghrelin pleiotropic roles. Apart from its well-described orexigenic effect, ghrelin impacts several other physiological functions, including memory and learning, sleep, metabolism, adipogenesis, immunity, reproduction, gastric motility, cardiac output, and bone formation, via centrally- and peripherally-mediated mechanisms (13-16), suggesting a significant role in organism aging (17, 18). Ghrelin levels and/or it is signaling are decreased in elderly individuals, which may be associated with age-related alterations in hypothalamic function and food intake, neuroendocrine outputs, metabolism, cardiovascular function, and bone remodeling (19-29). In addition, our group has previously shown that ghrelin stimulates autophagy, a proteostasis mechanism impaired in natural aging and HGPS (30, 31). Also, we and others showed that rapamycin, a known-autophagy stimulator, can decreases progerin and delay aging progression in HGPS cells (32, 33), therefore, ghrelin may act on similar pathways as rapamycin, exerting its beneficial effects through stimulation of autophagy and progerin clearance, ultimately rescuing the senescent phenotype of HGPS cells. Given its pleiotropic actions, we hypothesized that ghrelin delays or blocks the aging phenotype of HGPS and potentially increases lifespan. Here, using fibroblasts from HGPS patients and an animal model of HGPS, the *Lmna*^*G609G/G609G*^ mice, we report that ghrelin might delays premature aging of HGPS, and consequently improving longevity.

## Results

### Ghrelin increases autophagy and enhances progerin clearance in HGPS cells

One of the hallmarks of cellular aging in HGPS is loss of proteostasis and autophagy impairment, which could be one of the major promotors of progerin accumulation in HGPS cells (34). Using rodent neurons, we previously demonstrated that ghrelin activates autophagy (30, 31). Therefore, we hypothesized that ghrelin could promote progerin clearance through autophagy induction in fibroblasts from HGPS patients. Ghrelin treatment (6 hours) increased LC3B-II levels in HGPS cells (Fig. 1*A*). To investigate if this increase in LC3B-II was due to an inhibited autophagosome degradation rather than autophagosome formation, we evaluated the endogenous autophagic flux in the absence and presence of chloroquine, an inhibitor of autophagic degradation (35) (Fig. 1*B*), supporting that ghrelin stimulates autophagy in HGPS cells. Ghrelin also significantly decreased phospho-MTOR protein levels (Fig. 1*C*), indicating that the observed autophagy stimulation occurs by inhibiting MTOR in HGPS cells. This finding is consistent with our previous observations that rapamycin, positive control for autophagy induction, increased LC3B-II protein levels, a marker of autophagosome formation, and boosted autophagic flux through inhibiting MTOR (*Figure S1*, (33)). Concomitant with this upregulation in autophagy, ghrelin-treated HGPS cells showed lower progerin protein levels (Fig. 1*D*) compared to untreated HGPS cells. This effect was observed not only after 6 hours treatment, but also when HGPS cells were exposed to ghrelin for one week, treated every other day (Fig. 1*E*), suggesting that ghrelin leads to sustained progerin clearance. Rapamycin also decreased progerin levels (*Figure S1*, (33) and (32)), further supporting a role for autophagy in progerin clearance. Additionally, ghrelin-treated HGPS cells exhibited very low, or even absent, progerin immunoreactivity (Fig. 1*F*). Although progerin protein levels were decreased upon ghrelin treatment, its mRNA expression was not affected (*SI Appendix*, Fig. S1*A* and *B*). These results suggest that ghrelin promotes progerin clearance in HGPS cells.

**Figure 1.**
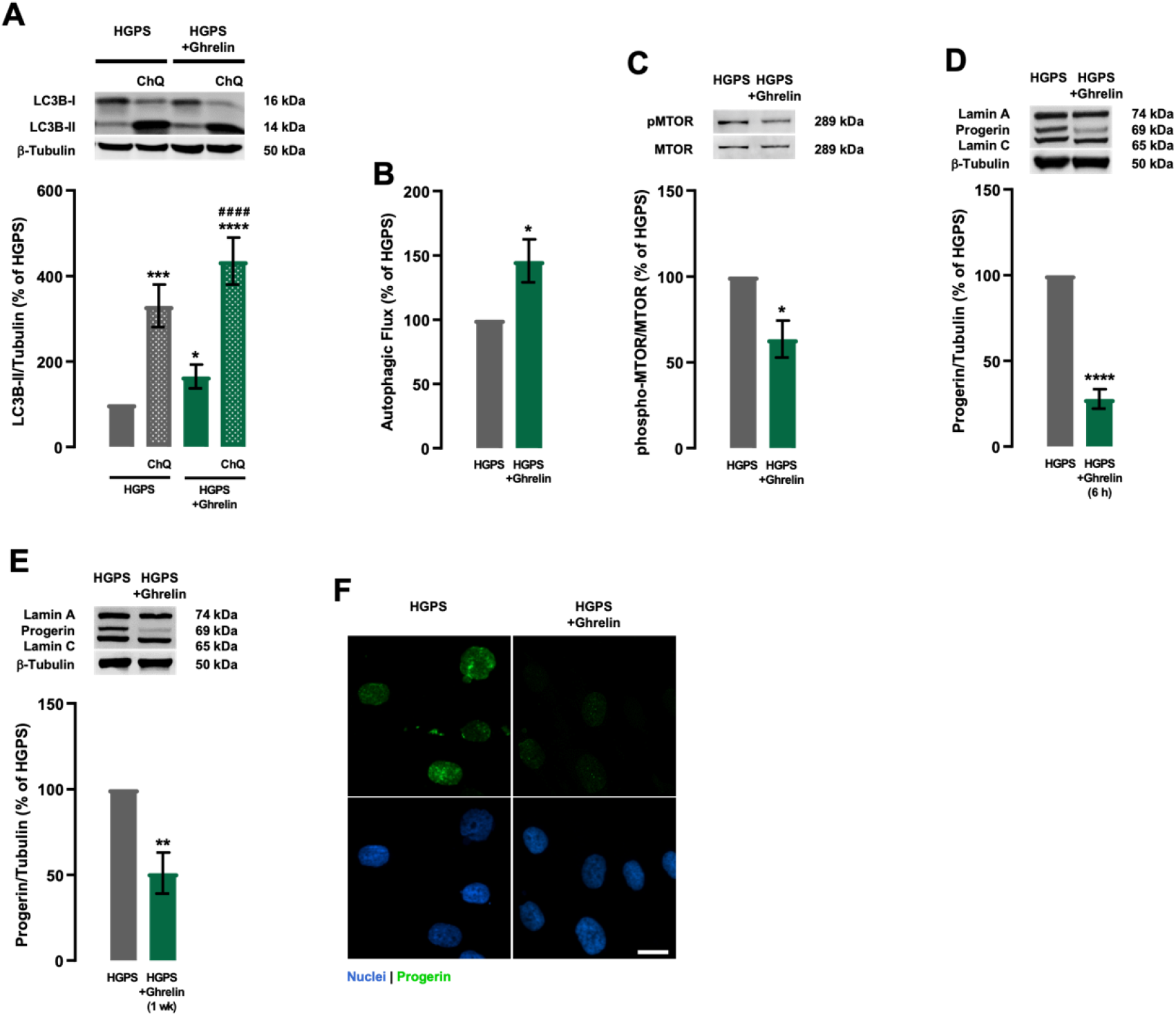
Ghrelin enhances autophagy and progerin clearance in HGPS fibroblasts. (*A-H*) HGPS fibroblasts were exposed to ghrelin (1 nM) for 6 hours (HGPS+Ghrelin) in the presence or absence of chloroquine (ChQ, 100 μM), a lysosomal degradation inhibitor (*A-D* and *G*), or for one week, treated every other day (*E* and *F*). HGPS-untreated cells were used as control (HGPS). Whole-cell extracts were assayed for LC3B (*A*) (*N*=4), pMTOR/tMTOR (*C*) (*N*=3), Lamin A/Progerin/Lamin C (*D* and *E*) (*N*=5) and β-Tubulin (loading control) immunoreactivity through Western blotting analysis. Representative Western blots for each protein are presented above each respective graph. Autophagic flux analysis in HGPS cells is shown (*B*) (*N*=4). Autophagic flux was determined in the presence of the lysosomal inhibitor chloroquine and expressed as “Autophagic flux” calculated by subtracting the densitometric value of LC3B-II-ChQ from those corresponding LC3B-II+ChQ values. (*F*) Ghrelin decreased progerin immunoreactivity. Cells were immunolabeled for progerin (top panels, green) and nuclei were stained with Hoechst 33342 (bottom panels, blue). Images are representative of three independent experiments. Scale bar, 10 μm. Data are expressed as the mean±SEM of at least three independent experiments and are expressed as a percentage of HGPS. ^*^*P*<0.05, ^**^*P*<0.01, ^***^*P*<0.001, and ^****^*P*<0.0001 significantly different compared to HGPS; ^####^*P*<0.0001, significantly different compared to HGPS+Ghrelin, as determined by analysis of variance, followed by Tukey’s multiple comparison test, or Student’s *t* test. HGPS = Hutchinson-Gilford progeria syndrome.

### Ghrelin rescues aberrant nuclear morphology and decreases DNA damage in HGPS cells

Given that ghrelin promotes progerin clearance, we further investigated ghrelin potential to rescue other cellular aging hallmarks of HGPS. After one week, ghrelin-treated HGPS cells exhibited a lower number of dysmorphic nuclei (Fig. 2*A* and *B*), one of several cellular defects in HGPS cells (7, 8). Furthermore, ghrelin decreased the frequency of nuclei with aberrant circularity (circularity <0.6; 24.62 ± 10.92 % in HGPS *vs* 3.29 ± 0.82 % in HGPS+Ghrelin) and an increased in the frequency of normal-shaped nuclei (circularity >0.8; 17.73 ± 2.32 % in HGPS vs 30.17 ± 6.81 % in HGPS+Ghrelin; Fig. 2*C*), suggesting that ghrelin, likely through the prevention of progerin accumulation, improves abnormal nuclear morphology of HGPS cells. Noteworthy, ghrelin also improved nuclear circularity and decreased the number of dysmorphic nuclei in primary cultures of fibroblasts from a healthy individual (*SI Appendix*, Fig. S2*A-C*). We next evaluated the effect of a 1-week ghrelin treatment on DNA damage through evaluation of the γ-H2AX *foci*, a DNA damage marker. As shown in Fig. 2*D*, ghrelin treatment decreased γ-H2AX immunoreactivity and the number of γ-H2AX *foci* in HGPS cells (Fig. 2*E*). These observations suggest that ghrelin decreases DNA damage in HGPS cells, probably by decreasing progerin accumulation.

**Figure 2.**
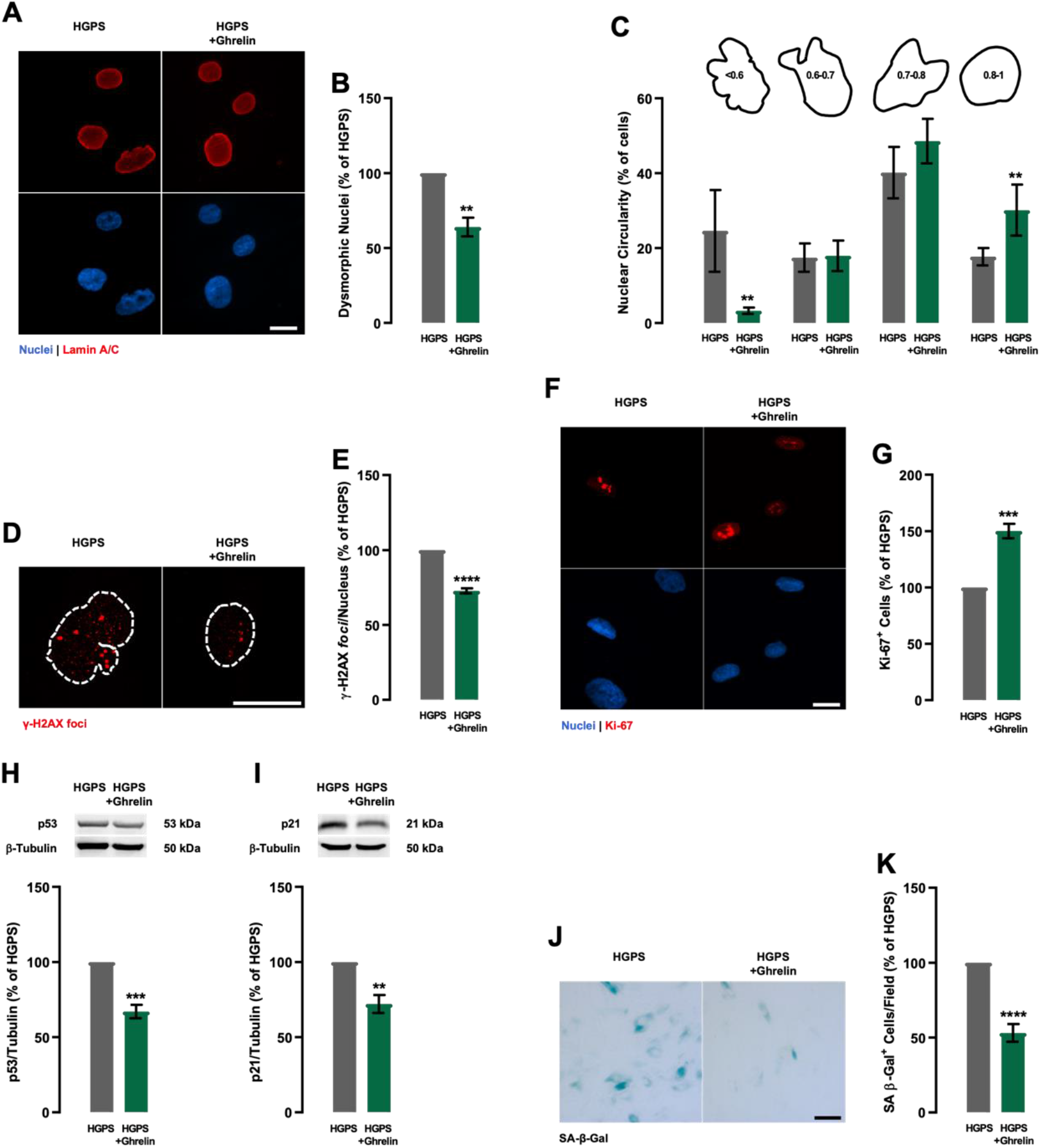
Ghrelin delays cellular senescence in HGPS fibroblasts. (*A-K*) HGPS fibroblasts were exposed to ghrelin (1 nM; HGPS+Ghrelin) for one week for every other day. HGPS-untreated cells were used as control (HGPS). (*A-C*) HGPS fibroblasts were immunolabeled for Lamin A/C (red, top panel) and nuclei were stained with Hoechst 33342 (blue, bottom panel) (*A*). Images are representative of four independent experiments. Scale bar, 10 μm. Quantification of the number of misshapen nuclei (*B*) and nuclear circularity (*C*) upon ghrelin treatment (*N*=4-6). For each condition, an equal number of nuclei (>400) were randomly analyzed. Circularity (defined as 4*π*area/perimeter^2) was measured using ImageJ. A circularity value equal to 1 corresponds to perfectly circular nuclei. (*D* and *E*) Ghrelin decreased γ-H2AX immunoreactivity. Cells were immunolabeled for γ-H2AX (red). Images are representative of three independent experiments. Scale bar, 10 μm. (*E*) Quantification of γ-H2AX *foci* number using ImageJ analysis and customized macros of three independent experiments; >200 cells analyzed). The average number of γ-H2AX *foci per* nucleus. (*F* and *G*) Ghrelin increases cell proliferation, as determined by Ki-67 immunoreactivity. (*F*) Cells were immunolabeled for Ki-67 (red, top panel) and nuclei were stained with Hoechst (blue, bottom panel). Images are representative of four independent experiments. Scale bar, 10 μm. (*G*) Quantification of the number of Ki-67-positive cells in HGPS and ghrelin-treated HGPS cells. (*H* and *I*) Whole-cell extracts were assayed for p53 (*H*) (*N*=4), p21 (*I*) (*N*=3), and β-Tubulin (loading control) immunoreactivity through Western blotting analysis. Representative Western blots for each protein are presented above each respective graph. (*J* and *K*) Ghrelin decreases cellular senescence, as determined by SA-β-Gal activity. (*J*) Images are representative of five independent experiments. Scale bar, 100 μm. (*K*) Quantification of SA-β-Gal-positive cells. Data are expressed as the mean±SEM, at least, three independent experiments, and are expressed as a percentage of HGPS. ^**^*P*<0.01,^**^*P*<0.001 and ^****^*P*<0.0001, significantly different from HGPS, as determined by Student’s *t* test. HGPS = Hutchinson-Gilford progeria syndrome.

### Ghrelin stimulates cell proliferation and delays cellular senescence in HGPS cells

Through assessment of Ki-67 immunoreactivity, we observed that 50 % of 1-week-ghrelin-treated HGPS cells showed positivity for this cell proliferation marker, compared to non-treated HGPS cells (Fig. 2*F* and *G*). This cell proliferation rescue was also observed in ghrelin-treated fibroblasts from a healthy individual (*SI Appendix*, Fig. S2*D* and *E*). The ghrelin-induced proliferative capacity of HGPS cells was accompanied by downregulation of p53 (Fig. 2*H*) and its downstream effector p21 (Fig. 2 *I*), well-known cell cycle repressors (36-38). Moreover, 1-week-ghrelin-treated HGPS cells also showed 50 % lower senescence-associated-β-Galactosidase (SA-β-Gal) activity (Fig. 2*J* and *H*). These results indicate that ghrelin may increase cell proliferation and postpone cellular senescence in HGPS cells by inhibiting the activation of the p53/p21 signaling pathway.

Overall, the results presented herein provide the basis for a model where ghrelin, via enhancement of autophagy, promotes progerin degradation and ameliorates several cellular defects typically associated with HGPS, including aberrant nuclear architecture, DNA damage, and cellular senescence.

### Ghrelin administration ameliorates age-dependent weight loss and extends the lifespan of *Lmna*^G609G/G609G^ mice

*Lmna*^G609G/G609G^ mice, generated by Osorio *et al*., carry the c.1827C>T;p.Gly609Gly mutation equivalent to the mutation found in HGPS patients (39). These mice recapitulate several features of HGPS, including progerin accumulation, reduced growth rate, low body weight, lipodystrophy, bone, and cardiovascular abnormalities, dysregulation of glucose and lipid metabolism, and shortened lifespan (39). Considering the beneficial effects of ghrelin observed in our *in vitro* model of HGPS (fibroblasts from HGPS patients), we next investigated if ghrelin could rescue the age-dependent phenotype in *Lmna*^G609G/G609G^ mice. For this purpose, we administered ghrelin daily to *Lmna*^G609G/G609G^ and *Lmna*^+/+^ littermates at 6-weeks of age, for 6-weeks (Fig. 3*A*). Recapitulating previous reports, *Lmna*^G609G/G609G^ mice exhibited a healthy appearance for 3-weeks after birth, after which they started to show a reduction in growth rate with progressive body weight loss (Fig. 3*B* and *C*). Ghrelin treatment increased body weight and attenuated body weight loss at later stages of *Lmna*^*G609G/G609G*^ mice, in contrast to vehicle-treated mice (Fig. 3*C*). The ghrelin-induced impact on body weight was independent of food intake (Fig. 3*D*). Additionally, ghrelin-treated *Lmna*^G609G/G609G^ mice showed an overall healthier appearance (Fig. 3*B*). Progeroid *Lmna*^G609G/G609G^ mice exhibited altered circulating plasma concentrations of several hormones and other biochemical markers, as well as low blood glucose levels (Fig. 3*E*). Ghrelin treatment induced a modest, but generalized, improvement in *Lmna*^G609G/G609G^ mice blood profile, especially in blood glucose and cholesterol levels (Fig. 3*E*). The organ size in progeroid mice was proportional to their reduced body weight (*SI Appendix*, Fig. S3*A*); however, with a significant decrease in liver and spleen weight (*SI Appendix*, Fig. S3*A*). Furthermore, these peripheral organs exhibited several histopathological alterations. Hepatocytes of progeroid *Lmna*^G609G/G609G^ mice liver were smaller and irregular, with reduced lipid vacuoles, compared to *Lmna*^+/+^ mice (*SI Appendix*, Fig. S3*B*, upper panel). Moreover, the lumen of sinusoidal capillaries is larger in progeroid *Lmna*^G609G/G609G^ mice than in *Lmna*^+/+^ (*SI Appendix*, Fig. S3*B*, upper panel). Ghrelin treatment did not affect the liver weight or histological structure of *Lmna*^G609G/G609G^ mice (*SI Appendix*, Fig. S3*A*). Masson’s trichrome staining was performed in liver sections to assess if there were alterations in collagen levels alterations induced by the accelerated aging process and to evaluate the effect of ghrelin on collagen deposition. We observed lower collagen staining in *Lmna*^G609G/G609G^ mouse liver, more prominent around the blood vessels, compared to *Lmna*^+/+^ mice (*SI Appendix*, Fig. S3*B*, bottom panel). Ghrelin treatment reestablished collagen levels in *Lmna*^G609G/G609G^ mice. Others have described a marked splenic involution in progeroid *Lmna*^G609G/G609G^ mice compared to wild-type controls, a feature that was associated with defective immune system of these animals (39). Similarly, we observed lighter spleens in *Lmna*^G609G/G609G^ mice than wild-type littermates. Ghrelin-treated *Lmna*^G609G/G609G^ mice showed a higher spleen weight compared to *Lmna*^+/+^ mice (*SI Appendix*, Fig. S3*A*). The histological structure of spleen was not significantly different between both genotypes, except for the white pulp area (*SI Appendix*, Fig. S3*C* and *D*) which was smaller in *Lmna*^G609G/G609G^ mice and rescued upon ghrelin treatment (Fig. 2*C* and *SI Appendix*, Fig. S3*D*). The reduced body weight of progeroid *Lmna*^G609G/G609G^ mice can be aggravated or associated with skeletal muscle atrophy. To investigate possible alterations in the skeletal muscle of *Lmna*^G609G/G609G^ mice and ghrelin treatment on this tissue, we measured the cross-sectional area of the muscular fibers. We did not observe significant differences in skeletal muscle histological structure in *Lmna*^G609G/G609G^ mice compared to *Lmna*^+/+^ mice, which showed polygonal or round muscle fibers with peripheral nuclei (*SI Appendix*, Fig. S3*E*). However, the cross-sectional area of skeletal muscle fibers was smaller in *Lmna*^G609G/G609G^ mice, when compared to *Lmna*^+/+^ mice (*SI Appendix*, Fig. *S3E* and *F*). Interestingly, ghrelin treatment rescued the cross-sectional area of skeletal muscle fibers in *Lmna*^G609G/G609G^ to the levels observed in *Lmna*^+/+^ mice (*SI Appendix*, Fig. *S3E* and *F*). These data demonstrate that ghrelin treatment ameliorates age-related alterations in peripheral organs, such as the liver, spleen, and skeletal muscle.

**Figure 3.**
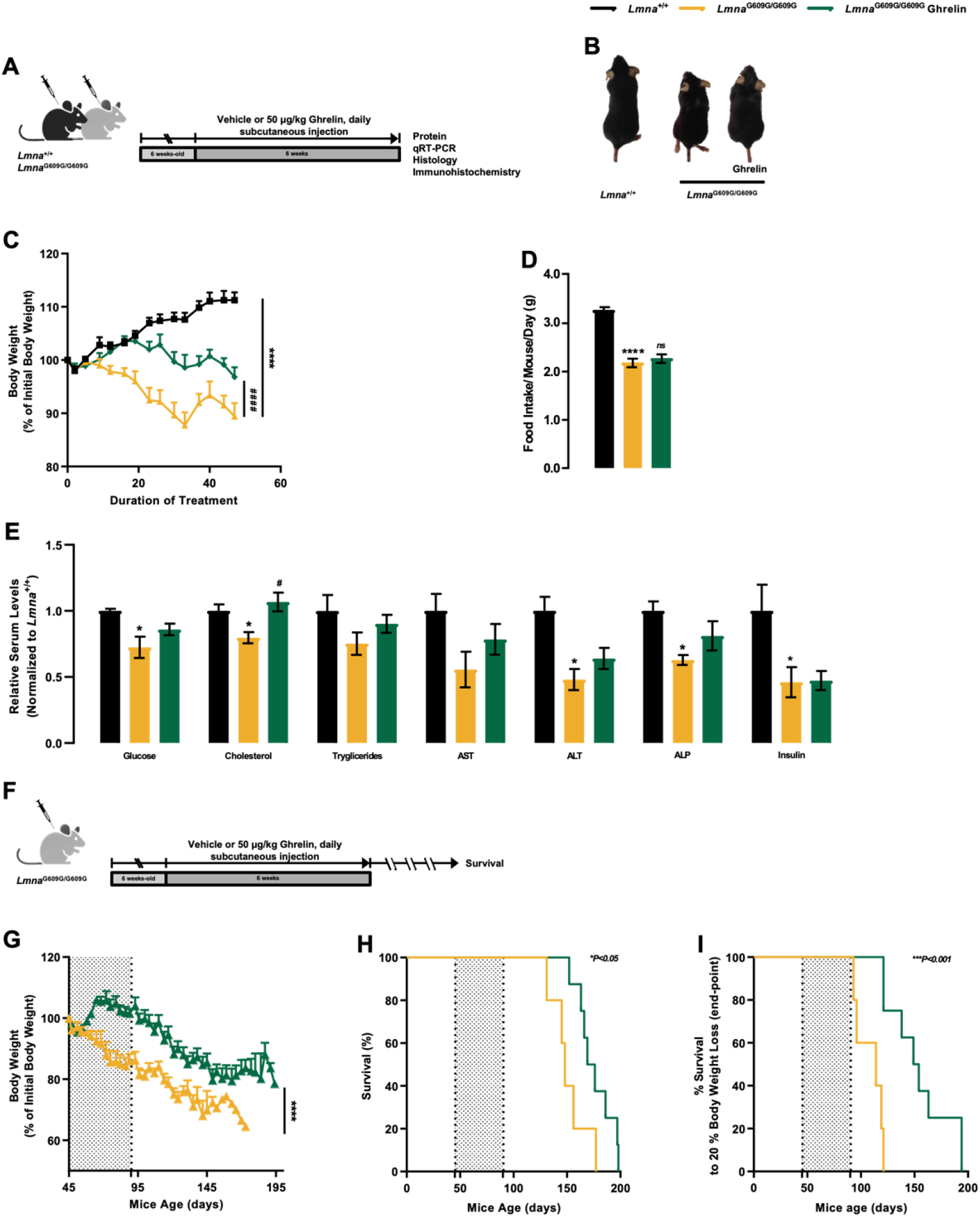
Ghrelin treatment directly affects *Lmna*^G609G/G609G^ phenotype improving the overall metabolic status without food intake changes, extending lifespan, and preventing age-associated phenotypes. (*A*) Schematic representation of the animal protocol. Effect of daily peripheral administration of ghrelin in *Lmna*^G609G/G609G^ mice premature aging phenotype. (*B*) Representative photographs of 3 months-old vehicle- or ghrelin-treated *Lmna*^+/+^ and *Lmna*^G609G/G609G^ mice. (*C*) Cumulative body weight gain of vehicle- and ghrelin-treated *Lmna*^+/+^ and *Lmna*^G609G/G609G^ mice, as the percentage of weight gain between the beginning and the end of the study. (*D*) Daily food intake (in grams) of vehicle- and ghrelin-treated *Lmna*^+/+^ and *Lmna*^G609G/G609G^ mice, expressed as g/day. (*E*) Serum concentration levels of glucose, cholesterol, triglycerides, aspartate transaminase (AST), alanine transaminase (ALT), alkaline phosphatase (ALP) and insulin of vehicle- and ghrelin-treated *Lmna*^+/+^ and *Lmna*^G609G/G609G^ mice. Serum concentrations were normalized to the mean of *Lmna*^+/+^. Data are expressed as the mean±SEM. *N*=4-19 *per* group. ^*^*P*<0.05 and ^****^*P*<0.0001, significantly different from *Lmna*^+/+^ mice; ^#^*P*<0.05 and ^###^*P*<0.001, significantly different compared to *Lmna*^G609G/G609G^ mice, as determined by one analysis of variance, followed Tukey’s multiple comparison test. (*F*) Schematic representation of the animal protocol. Effect of daily peripheral administration of ghrelin in *Lmna*^G609G/G609G^ mice health decline. (*G*) Cumulative body weight gain of vehicle- and ghrelin-treated *Lmna*^G609G/G609G^ mice, as the percentage of weight gain between the beginning and the end of the study. (*H*) Kaplan-Meier survival plots for vehicle- and ghrelin-treated *Lmna*^G609G/G609G^ mice. (*I*) Ghrelin-treated *Lmna*^G609G/G609G^ mice show a 37 % increased median age at end-point, compared to vehicle-treated *Lmna*^G609G/G609G^ (152 days *versus* 114 days respective median age at end-point; based on mice being terminated upon reaching a 20 % body weight loss); and more than 73 days between the longest-lived (20 % body weight loss) ghrelin-treated *Lmna*^G609G/G609G^ and the longest-lived vehicle-treated *Lmna*^G609G/G609G^ mouse. Data are expressed as the mean±SEM. *N*=5-8 *per* group. ^*^*P*<0.05 and ^***^*P*<0.001, significantly different compared to *Lmna*^G609G/G609G^ mice, as determined by Student’s *t* test and Log-rank/Mantel-Cox test; Chi-Square 5.19 and 11.11 for graphs (*H*) and (*I*), respectively.

As previously described, *Lmna*^G609G/G609G^ mice have a dramatically shortened life expectancy compared to *Lmna*^+/+^ mice (39). Given the improved weight loss and growth retardation observed in progeroid mice treated with ghrelin, we hypothesized that ghrelin might have a beneficial effect on the lifespan of *Lmna*^G609G/G609G^ mice. Ghrelin-treated *Lmna*^G609G/G609G^ mice showed significant body weight improvement during their lifespan and lower weight loss at later stages of the disease (Fig. 3*G*) exhibiting significant healthier aspect than vehicle-treated mice. Moreover, ghrelin treatment extended the lifespan of progeroid *Lmna*^G609G/G609G^ mice (Fig. 3 *H* and *I*). The mean survival time of ghrelin-treated *Lmna*^G609G/G609G^ mice increased from 148 to 173 days and the maximum survival time increased from 177 to 198 days, representing a ∼22 % increase in the Kaplan-Meier area under the curve (AUC) (Fig. 3*H*). The increase in progeroid mice lifespan may relate with delayed body weight loss upon ghrelin treatment. In fact, when the Kaplan-Meier AUC is adjusted for 20 % body weight loss, the beneficial effects of ghrelin were even more noticeable between ghrelin-treated *Lmna*^G609G/G609G^ mice *versus* the vehicle-treated *Lmna*^G609G/G609G^ mice (Fig. 3*I*). Indeed, we observed a 73-day gap between the longest-lived *Lmna*^G609G/G609G^ ghrelin-treated mouse, based on 20 % body weight loss, and the longest-lived *Lmna*^G609G/G609G^ vehicle-treated mouse (Fig. 3*I*). Overall, our results show that ghrelin treatment promotes considerable benefits to the healthspan of progeroid mice, as shown by the effects on age-dependent body weight loss in *Lmna*^G609G/G609G^ mice, extending lifespan.

### Ghrelin ameliorates aging markers in the aorta and heart of *Lmna*^G609G/G609G^ mice

The cardiovascular system is severely affected in HGPS patients, with myocardial infarction being the most common cause of death (2). *Lmna*^G609G/G609G^ mice recapitulate, to some degree, the abnormalities described in these patients, as they exhibit a depletion of vascular smooth muscle cells (VSMCs) in the aortic arch, which may underlie the cardiovascular system dysfunction (39). We observed a slight but nonsignificant decrease in medial aortic thickness (Fig. 4*A* (upper panel) and *B*) and a decreased cell number in this structure of *Lmna*^G609G/G609G^ mice (Fig. 4*A* (bottom panel) and *C*), compared with *Lmna*^+/+^ mice. Ghrelin treatment promoted a slight increase in both aortic thickness (Fig. 4*A* (upper panel) and *B*) and cell density (Fig. 4*A* (bottom panel) and *C*). Progerin induces a dramatic effect on VSMCs, leading to decreased cell viability and increased arterial stiffness (40). We observed a decrease in alpha-smooth muscle actin (α-SMA) immunoreactivity in the aortic wall of *Lmna*^G609G/G609G^ mice compared to *Lmna*^+/+^ mice (Fig. 4*D*), which correlates with the loss of cellularity (Fig. 4*A* and *C*). Ghrelin treatment slightly increased α-SMA immunoreactivity in *Lmna*^G609G/G609G^ mice (Fig. 4*D*). We also observed a strong progerin immunoreactivity encircling the nuclei of *Lmna*^G609G/G609G^ mice (Fig. 4*E*)., that was decreased upon ghrelin treatment (Fig. 4*E* and *F*). Additionally, no histological changes were observed in the heart, between the two genotypes, nor any effect deriving from ghrelin treatment (data not shown). Nevertheless, the hearts of *Lmna*^G609G/G609G^ mice treated with ghrelin showed lower progerin levels (Fig. 4*G*). These results show that ghrelin prevents VSMC loss and improves aortic structure in *Lmna*^G609G/G609G^ mice, which may be related to the ghrelin-induced decrease in progerin levels. Ghrelin might also have putative beneficial effects on heart functions by lowering progerin levels.

**Figure 4.**
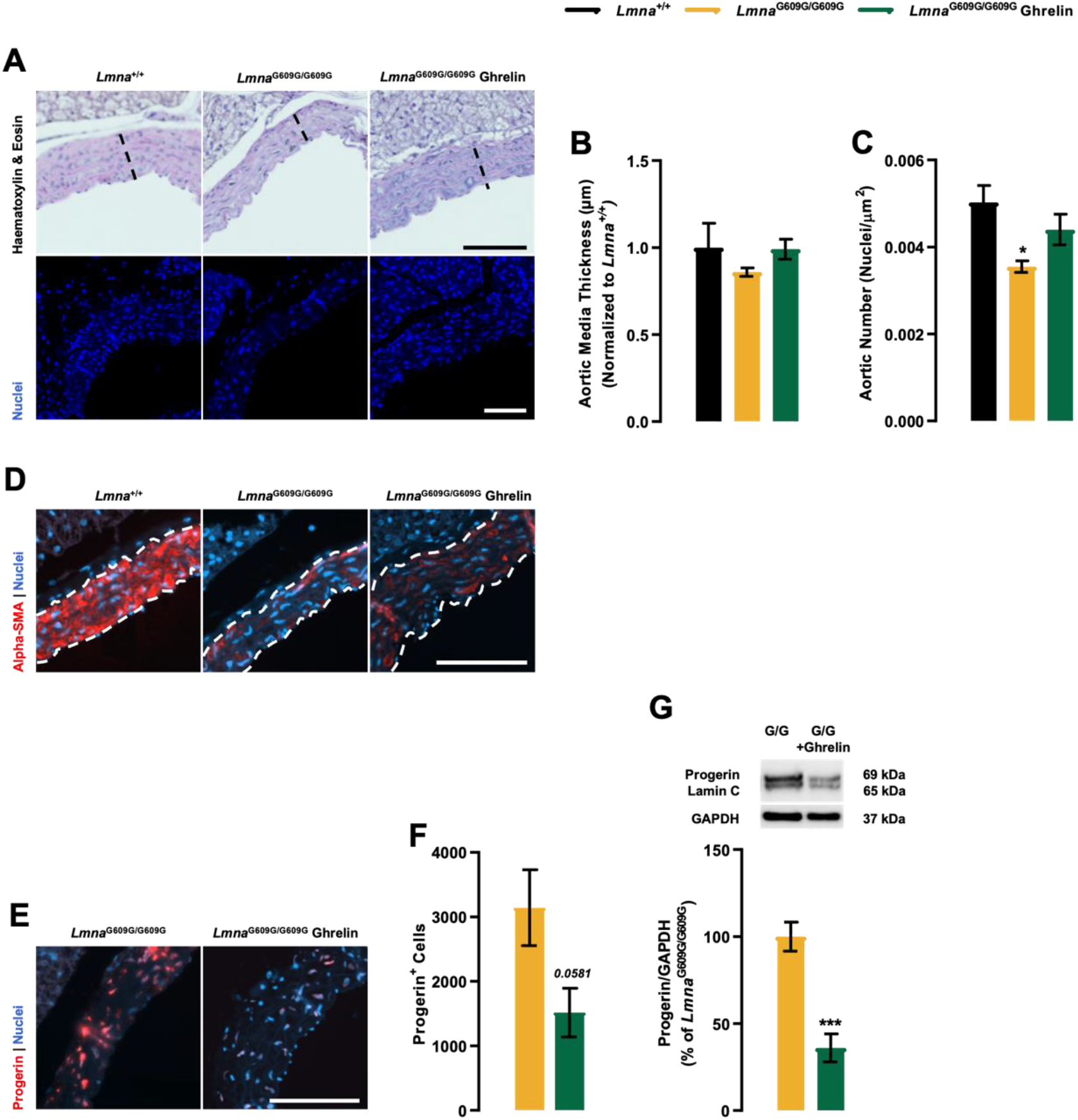
Ghrelin ameliorates cardiac related-pathology of *Lmna*^G609G/G609G^ mice. (*A*) Representative images of Hematoxylin-eosin-(top panel) and Hoechst 33342-(blue, bottom panel) stained cross-sections of the aorta of vehicle- and ghrelin-treated *Lmna*^+/+^ and *Lmna*^G609G/G609G^ mice. Scale bar, 100 μm. (*B* and *C*) Quantification of aortic media wall thickness, expressed in μm (*B*), and aortic wall nuclei number, expressed as aortic wall nuclei number/mm^2^ (*C*) in the aorta of vehicle- and ghrelin-treated *Lmna*^+/+^ and *Lmna*^G609G/G609G^ mice. (*D*) Representative images of aorta sections of vehicle- and ghrelin-treated *Lmna*^+/+^ and *Lmna*^G609G/G609G^ mice immunolabeled for alfa-smooth muscle actin, α-SMA, (red). Nuclei are stained with Hoechst 33342 (blue). Scale bar, 100 μm. (*E*) Representative images of the aorta of vehicle- and ghrelin-treated *Lmna*^+/+^ and *Lmna*^G609G/G609G^ mice immunolabeled for progerin (red). Nuclei are stained with Hoechst 33342 (blue). Scale bar, 100 μm. (*F*) Quantification of progerin-positive cells in the aorta expressed in cell/mm^2^. (*G*) Heart whole protein lysates were assayed for Lamin A/Progerin/Lamin C and GAPDH (loading control) immunoreactivity through Western blotting analysis. The results are expressed as a percentage of *Lmna*^G609G/G609G^ mice. Representative Western blots for each protein are presented in the graph. Data are expressed as the mean±SEM. *N*=4-6 *per* group. ^*^*P*<0.05, significantly different from *Lmna*^+/+^ mice, ^***^*P*<0.001, significantly different compared to *Lmna*^G609G/G609G^ mice, as determined by analysis of variance, followed Tukey’s multiple comparison test or Student’s *t* test.

### Ghrelin ameliorates skin thinning and subcutaneous adipose tissue atrophy in *Lmna*^G609G/G609G^ mice

In HGPS patients, skin presents relevant alterations: hyper- and hypopigmentation, gradual loss of the subcutaneous fat layer, which makes the skin extremely thin, prominent blood vessels, particularly those on the face and scalp, and loss of hair (2, 41). Others described some of these skin features in *Lmna*^G609G/G609G^ mice, namely the decrease in the subcutaneous fat layer and hair follicle attrition (39). Using histomorphometry analysis of dorsal skin (Fig. 5*A*, upper panel) we observed that *Lmna*^G609G/G609G^ mice exhibited thinning of the epidermis compared to *Lmna*^+/+^, a feature rescued with ghrelin treatment (Fig. 5*B*). Furthermore, progeroid *Lmna*^G609G/G609G^ mice showed a thinner dermis when compared to *Lmna*^+/+^ mice, however, ghrelin treatment had no impact on this structure (Fig. 5*C*). *Lmna*^G609G/G609G^ mice showed severe thinning and atrophy of the subcutaneous fat layer, the hypodermis, (Fig. 5*D*). Ghrelin-treated *Lmna*^G609G/G609G^ mice displayed a thicker subcutaneous adipose layer, suggesting that ghrelin ameliorates the lipodystrophic phenotype of progeroid mice (Fig. 5*D*). The degradation of collagen fibers is one of the major age-associated alterations in the skin. In comparison to *Lmna*^+/+^ mice, the skin of *Lmna*^G609G/G609G^ mice contained lower levels of collagen (Fig. 5*A*, bottom panel), and the collagen bundles appear to be organized in a similar fashion with open spaces between the collagen fibers. However, treatment with ghrelin increased collagen deposition in *Lmna*^G609G/G609G^ mice, suggesting that ghrelin may promote collagen synthesis or remodeling (Fig. 5*A*). To evaluate the proliferative capacity of skin cells, we used Ki-67 immunostaining and observed a lower number of Ki-67-positive cells in the skin of the progeroid mice (Fig. 5*E*, upper panel); however, the Ki-67-positive cells were arranged in niches, especially in the hair follicles bulb and in some regions of the epidermis (Fig. 5*F*). Ghrelin treatment increased the number of Ki-67-positive cells in these areas, indicating that ghrelin promotes proliferation in the epidermis of the *Lmna*^G609G/G609G^ mice skin (Fig. 5*F*). We also evaluated skin structure by examining the immunoreactivity of Keratin-1 (KRT1), a critical marker of skin integrity (Fig. 5*F*, bottom panel). The skin of *Lmna*^G609G/G609G^ mice exhibited lower KRT1 immunoreactivity compared to *Lmna*^+/+^ mice (Fig. 5*G*), however, this was improved with ghrelin treatment, suggesting that ghrelin can improve skin structure and integrity. Additionally, we observed strong progerin immunoreactivity surrounding the nuclei of skin cells in *Lmna*^G609G/G609G^ mice. Ghrelin treatment decreased progerin immunoreactivity and/or the density of progerin-positive cells (Fig. 5*I*). Overall, these findings indicate that ghrelin can ameliorate age-related changes in the skin of *Lmna*^G609G/G609G^ mice.

**Figure 5.**
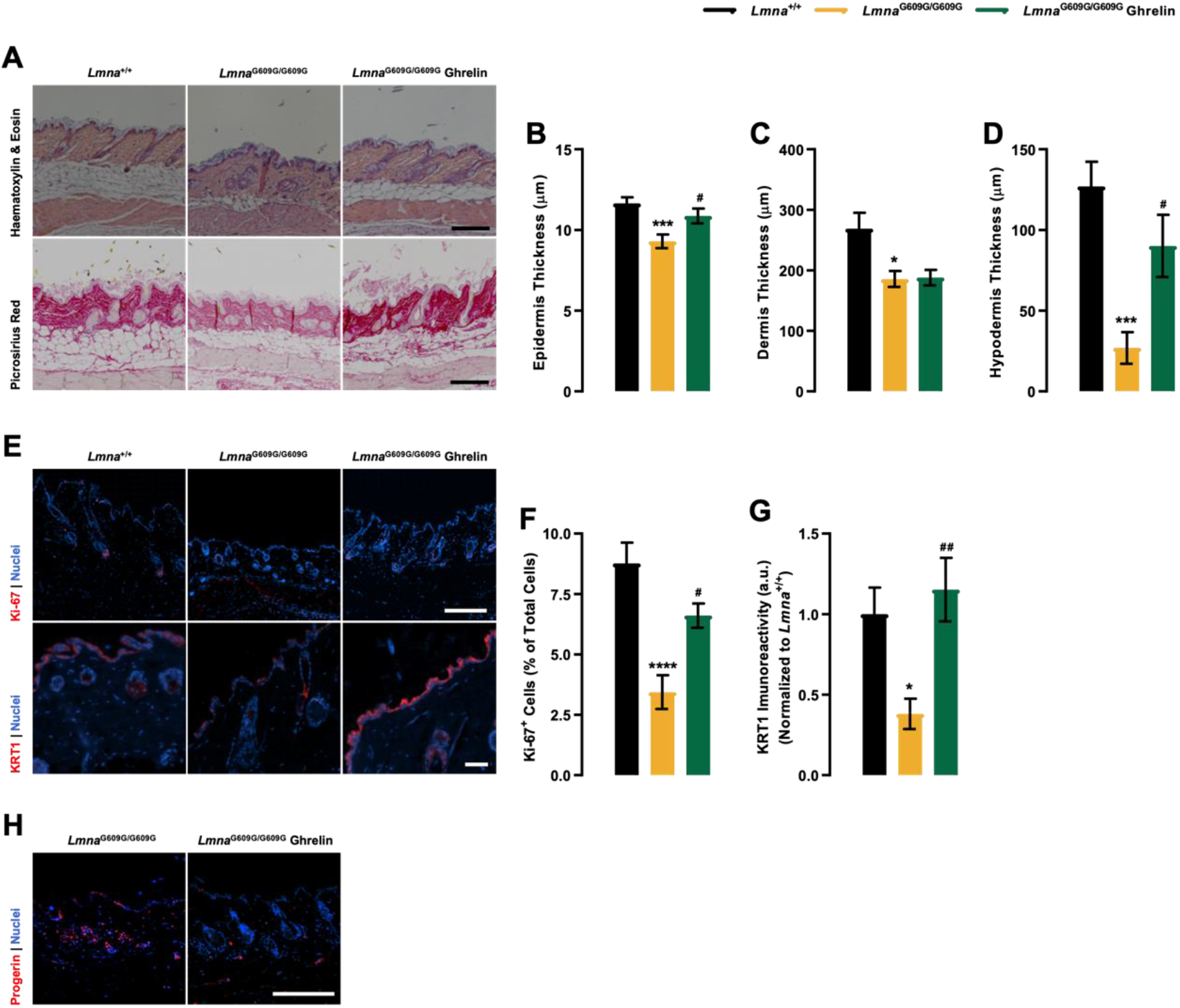
Ghrelin delays skin age-related alterations in the skin of *Lmna*^G609G/G609G^ mice. (*A*) Representative images of Hematoxylin-eosin-stained (top panel) and Picro-Sirius-Red-stained (bottom panel) sections of dorsal skin of vehicle- and ghrelin-treated *Lmna*^+/+^ and *Lmna*^G609G/G609G^ mice. Scale bar, 100 μm. (*B-D*) Quantification of the epidermis (*B*), dermis (*C*) and subcutaneous falt layer thickness (*D*) (expressed in μm), of the dorsal skin of vehicle- and ghrelin-treated *Lmna*^+/+^ and *Lmna*^G609G/G609G^ mice. (*E*) Representative images of dorsal skin of vehicle- and ghrelin-treated *Lmna*^+/+^ and *Lmna*^G609G/G609G^ mice immunolabeled for Ki-67 (red, top panel) and/or Keratin-1 (red, bottom panel). Nuclei are stained with Hoechst 33342 (blue). Scale bar, 100 μm. (*F*) Quantification of Ki-67-positive cells in the epidermal layer of the skin, expressed in % of total cells. (*G*) Quantification of KRT1 immunoreactivity in the epidermal layer of the skin, expressed in a.u, normalized to the mean of *Lmna*^+/+^. (*H*) Representative images of dorsal skin sections of vehicle- and ghrelin-treated *Lmna*^+/+^ and *Lmna*^G609G/G609G^ mice immunolabeled for progerin (red). Nuclei are stained with Hoechst 33342 (blue). Scale bar, 100 μm. Data are expressed as the mean±SEM. *N*=8-14 *per* group. ^*^*P*<0.05, ^***^*P*<0.001 and ^****^*P*<0.0001, significantly different from *Lmna*^+/+^; ^#^*P*<0.05 and ^##^*P*<0.01, significantly different compared to *Lmna*^G609G/G609G^ mice, as determined by analysis of variance, followed Tukey’s multiple comparison test.

### Ghrelin rescues the lipodystrophic phenotype of *Lmna*^G609G/G609G^ mice by reverting progerin-affected genes in the adipogenic network

*LMNA* mutations are associated with lipodystrophic features, which combine generalized or partial fat atrophy and metabolic alterations that could result from altered adipocyte differentiation or altered fat structure (42, 43), features also observed in *Lmna*^G609G/G609G^ mice (39). Here, we observed a significantly lower gonadal white adipose tissue (WAT) weight in *Lmna*^G609G/G609G^ mice compared with *Lmna*^+/+^ mice, which was reversed upon ghrelin treatment (Fig. 6*A*). Histomorphometry analysis of gonadal fat also revealed significantly reduced adipocyte area in *Lmna*^G609G/G609G^ mice (Fig. 6*B* and *C*). Concomitant with a WAT increase, ghrelin treatment induced an increase in adipocyte cross-sectional area, as well as a decrease in adipocyte density, due to the increased adipocyte size in *Lmna*^G609G/G609G^ mice (Fig. 6*C* and *D*), rescuing WAT fibrosis in progeroid mice (Fig. 6*B*). Progerin accumulation results in metabolic dysfunction, one crucial hallmark of the progeria phenotype, and ghrelin treatment decreased progerin protein levels in *Lmna*^G609G/G609G^ mice (Fig. 6*E*). Regarding brown adipose tissue (BAT), *Lmna*^G609G/G609G^ mice showed a lower quantity of BAT, when compared to *Lmna*^+/+^ (*SI Appendix*, Fig. S3*G*). with smaller adipocytes and with fewer lipid inclusions (*SI Appendix*, Fig. S3*H*). Ghrelin treatment rescued not only *Lmna*^G609G/G609G^ BAT weight (Fig. 6*G*) but also BAT architecture (*SI Appendix*, Fig. S3*H*). As in WAT, BAT of ghrelin-treated *Lmna*^G609G/G609G^ mice showed lower progerin levels compared to untreated-progeroid mice (*SI Appendix*, Fig. S3*I*). The structural abnormalities observed in the WAT of *Lmna*^G609G/G609G^ mice may be caused by an impairment of the adipogenic process, which may significantly impact the endocrine function of this organ and consequently regulate body weight and metabolism. We next investigated how ghrelin impacts WAT function by analyzing the levels of relevant hormones produced by WAT -leptin, adiponectin, and resistin. Consistent with their lipodystrophic phenotype, *Lmna*^G609G/G609G^ mice exhibited lower serum or gene expression of these hormones, a feature that was counteracted through ghrelin treatment (Fig. 6*F* and *G*).

**Figure 6.**
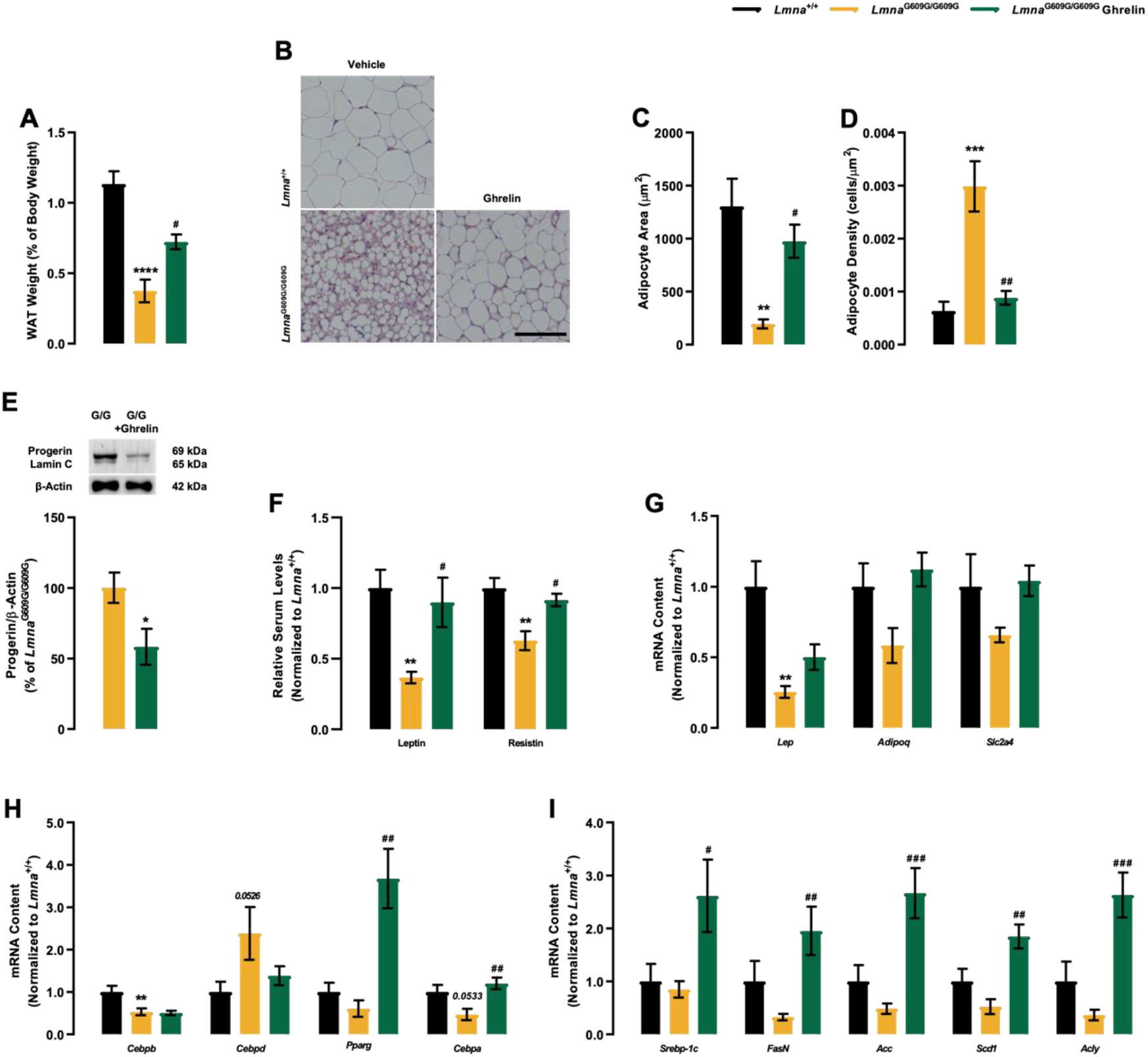
Ghrelin alleviates pathological changes in fat distribution and reverts the progerin-impacted transcription of core adipogenic regulators during adipocyte differentiation of *Lmna*^G609G/G609G^. (*A*) Size of the gonadal white adipose tissue (WAT), expressed as a percentage of % of body weight in 3 months-old vehicle- and ghrelin-treated *Lmna*^+/+^ and *Lmna*^G609G/G609G^ mice. (*B*) Representative images of Hematoxylin-Eosin-stained sections of WAT of vehicle- and ghrelin-treated *Lmna*^+/+^ and *Lmna*^G609G/G609G^ mice. Scale bar, 100 μm. (*C*) Quantification of adipocyte area (μm^2^) of vehicle- and ghrelin-treated *Lmna*^+/+^, and *Lmna*^G609G/G609G^ mice. (*D*) Quantification of adipocyte density (cells/μm^2^) of vehicle- and ghrelin-treated *Lmna*^+/+^ and *Lmna*^G609G/G609G^ mice. (*E*) WAT protein lysates were assayed for Lamin A/Progerin/Lamin C and β-Actin (loading control) immunoreactivity through Western blotting analysis. The results are expressed as a percentage of *Lmna*^G609G/G609G^ mice. Representative Western blots for each protein are presented in the graph. (*F*) Serum concentration levels of leptin and resistin of vehicle- and ghrelin-treated *Lmna*^+/+^ and *Lmna*^G609G/G609G^ mice. Serum concentrations were normalized to the mean of *Lmna*^+/+^. (*G-I*) Quantitative polymerase chain reaction analysis mRNA levels of (*G*) *Lep, Adipoq* and *Slc2a4* (*H*) WAT adipogenic differentiation genes (*Cebpb, Cebpd* (early differentiation regulators), *Pparγ* and *Cebpab* (late differentiation regulators)), (*I*) lipogenic genes (*Srebp-1c, FasN, Acc, Scd1* and *Acly*), fatty acid β-oxidation (*Mcad* and *Cpt1*) and gluconeogenic gene (*Pepck*), mRNA contents were normalized to the mean of *Lmna*^+/+^ mice. Data are expressed as the mean±SEM. *N*=5-12 *per* group. ^*^*P*<0.05, ^**^*P*<0.01 and ^****^*P*<0.0001, significantly different from *Lmna*^+/+^ or *Lmna*^G609G/G609G^ mice; ^#^*P*<0.05, ^##^*P*<0.01 and ^##^*P*<0.01, significantly different compared to *Lmna*^G609G/G609G^ mice, as determined by analysis of variance, followed Tukey’s multiple comparison test or Student’s *t* test.

The adipocyte lifecycle is activated by an initial transient increase of CCAAT-enhancer-binding proteins (C/EBP) isoforms β and δ, which, in response to adipogenic signals, precede the induction of peroxisome proliferator-activated receptor-gamma (PPARγ) and C/EBPα (44). In the WAT of *Lmna*^G609G/G609G^ mice, *Cebpb, Ppar*γ, and *Cebpa* were downregulated while *Cebpd* was upregulated compared to *Lmna*^+/+^ mice. Ghrelin treatment had no impact on *Cebpb* expression, but rescued the expression of *Cebpd, Ppar*γ, and *Cebpa* (Fig. 7*H*). Moreover, several enzymes involved in *de novo* lipogenesis (*Srebp-1c, FasN, Acc, Scd1* and *Acly*), fatty acid oxidation (*Mcad* and *Cpt1*), and *gluconeogenesis* (*Pepck*), showed decreased expression in *Lmna*^G609G/G609G^ mice, which was reverted with ghrelin treatment (Fig. 7*I*). These data indicate that mutations in *Lmna* resulted in adipogenic and metabolic defects. Ghrelin treatment promotes WAT differentiation and maturation and improves WAT function in *Lmna*^G609G/G609G^ mice concomitant with progerin clearance, which could counteract the lipodystrophy observed in this mouse model.

## Discussion

HGPS is a rare and lethal genetic condition that results in premature and accelerated aging. This disorder arises from a spontaneous point mutation in the *LMNA* gene, which gives rise to a defective prelamin A form, known as progerin (5, 6). HGPS represents a paradigm for translational medicine in aging. The identification of novel therapeutic agents for this disease is crucial and provides promising strategies for delaying the aging process. Numerous treatments have been suggested to improve the HGPS condition, such as rapamycin, sulforaphane, and MG132, which are known to enhance proteostasis and progerin clearance, leading to an improvement of aging-related defects in HGPS cells (32, 34, 45). Although promising for treating HGPS *in vivo*, cautious strategies are still needed before translation into clinical trials.

Rescue of autophagy impairment delays the aging progression of aging in HGPS (32, 34, 45); therefore ghrelin, a hormone that stimulate autophagy and progerin clearance, could benefit HGPS phenotype. Moreover, ghrelin might also benefit other HGPS features through its known effects on body weight, white adipose tissue, metabolism, cardiovascular function, and bone mass regulation (46). Here we show, for the first time, that ghrelin enhances progerin clearance and delays cellular senescence in HGPS cells. Importantly, we demonstrated that ghrelin treatment ameliorates the aging phenotype and extends the lifespan of HGPS mice. These findings suggest that ghrelin could be a promising therapeutic candidate for HGPS and other age-related diseases. First, we found that ghrelin reduces the accumulation of progerin in HGPS cells by enhancing autophagic flux. This result aligns with our previous reports in rodent neurons showing that ghrelin stimulates autophagy (30, 31). One of the most notable characteristics of HGPS cells is their abnormal nuclear shape, which results from the accumulation of progerin in the nuclear membrane (8, 32). Ghrelin improves nuclear architecture, decreasing the number of deformed nuclei, which can result from the progerin clearance from the nuclear envelope. Consequently, the normal lamin A protein could interact with the nuclear envelope, devoid of progerin, and restore the normal nuclear scaffold, ensuring the stability and integrity of the nucleus of these cells. Ghrelin also reduces DNA damage in HGPS cells; through the decrease of progerin levels, ghrelin may restore the connection between the lamins and DNA which may lead to a reduction in DNA damage and ultimately prevent genomic instability (47, 48). Ghrelin also improves the proliferative cell capacity, through decreased activation of the p53/p21 pathway. Chronic activation of p53 due to permanent DNA damage is a common denominator between cell senescence and DNA damage in HGPS patients (49, 50). Ghrelin treatment led to reduced progerin levels and nuclear abnormalities, which triggered the activation of checkpoint response to nuclear abnormalities. This inhibited the stress signaling pathways activated by p53 and enhanced the DNA damage response machinery, thereby promoting reactivation of the cell cycle (38, 49, 51).

Second, we have demonstrated that treating HGPS mice with ghrelin ameliorates the main age-related histopathological alterations of mesenchymal tissues (52), such as lipodystrophy, skin thinning, liver fibrosis, muscle atrophy, and aortic thinning, ultimately attenuating the age-related functional decline, and extending the health and lifespan of progeria mice. Current evidence suggests that metabolic and endocrine alterations contribute to the HGPS phenotype (46). Adipose tissue is an endocrine and metabolic organ that plays an important role in energy homeostasis (53). It also regulates several body functions, such as blood pressure, reproduction, angiogenesis, and immune response (54-56). The progeria mouse model used in this study -*Lmna*^G609G/G609G^ mice -successfully recapitulates the patient adipose tissue phenotype (57-59), showing smaller adipocytes that may suggest an impaired capacity to store lipids and may represent one of the causes of lipodystrophy in these animals. One of the major findings of our study is that ghrelin treatment can rescue the structural abnormalities found in the white adipose tissue (WAT) adipocytes and, consequently, WAT loss (severe lipodystrophy) in HGPS mice. This beneficial effect of ghrelin may be explained not only by the well-described ghrelin action on adiposity and energy expenditure inhibition (60, 61), but also from its direct action on WAT-promoting adipocyte differentiation, maturation, and lipid accumulation. Capanni and colleagues reported an *in vivo* interaction between prelamin A and SREBP1, a transcription factor upstream of PPARγ and essential for adipocyte differentiation. This interaction resulted in the sequestration of SREBP1 from the nuclear interior to the periphery, leading to the downregulation of its target genes (62). It is possible that progerin, known to sequester several transcriptional factors at the nuclear periphery through a similar mechanism, may also promote similar changes (63, 64). Based on our findings, we hypothesize that the structural changes we observed in the WAT of *Lmna*^G609G/G609G^ mice may be caused by an impairment of the adipogenic process. Adipogenesis is controlled by a cascade of fat cell-related transcriptional factors (44). We found that C/EBPβ, an early adipogenic gene, as well as PPARγ and C/EBPα, essential for adipogenic differentiation and maturation, were downregulated in *Lmna*^G6009G/G609G^ mice. Furthermore, ghrelin treatment rescued adipocyte maturation, related to decreased progerin levels and improved adipocyte architecture and function. Ghrelin treatment also restored leptin levels, an important hormone for WAT function (65), and the expression of several key regulators of lipogenesis, fatty acid oxidation, and glucose homeostasis in WAT from *Lmna*^G609G/G609G^ mice. Ghrelin treatment also improved the serum levels of cholesterol, triglycerides, and glucose, improving the overall metabolic status of the mice. However, the increased glucose levels observed may be related to the inhibition of insulin secretion, which has been described because of ghrelin (66). The effect of ghrelin on blood glucose levels could be a protective mechanism, as it may minimize hypoglycemia by decreasing glucose uptake by different tissues during fasting periods when glucose levels are low. Since progeroid mice show hypoglycemia (39), which could aggravate the cardiovascular abnormalities observed in these animals (67, 68), the effect of ghrelin on blood glucose levels could have a beneficial impact on cardiovascular functions and minimize the progression of the disease. The skin of HGPS patients is also severely affected by disease progression, resulting in thinner skin and contributing to the lipodystrophic phenotype (41). Skin alterations have also been reported in *Lmna*^G609G/G609G^ mice (39), including loss of the subcutaneous fat layer and wear of hair follicles. We also found tha*t Lmna*^G609G/G609G^ mice showed an overall thinning of the epidermis, dermis, and hypodermis due to decreased cell proliferative capacity. However, the subcutaneous fat layer changes are more pronounced, probably related to severe lipodystrophy in these animals (39). With the normal aging process, the skin also becomes thinner as the number of epidermal cells decreases, the vascularity and cellularity of the dermis decrease, and the subcutaneous fat in the hypodermis falls (69). These processes are exacerbated in HGPS mice, accelerating the aging phenotype. We found that ghrelin treatment restored epidermal layer thinning in *Lmna*^G609G/G609G^ mice, which may be triggered by increased cell proliferation. One hypothesis to explain the observed beneficial effects of ghrelin treatment is that it restored epidermal layer thinning in *Lmna*^G609G/G609G^ mice, which may result from increased cell proliferation. Ghrelin may also induce the proliferation of pre-existing keratinocytes, considering its well-established role as a cell proliferation stimulator (18, 70-74). Another possible mechanism underlying this effect is the increased stem cell proliferation and differentiation, which has already been described in diverse models (75, 76). Age-associated skin changes also include the decrease in the extracellular matrix components, such as collagen (69). Ghrelin treatment attenuated collagen loss in progeroid mice, suggesting increased collagen synthesis and turnover. However, further studies are required to explore collagen and extracellular matrix alterations in the skin of these mice, given that they are also related to cell proliferative capacity (77-79).

Overall, we conclude that, ghrelin improved the healthspan and extended the lifespan of *Lmna*^G609G/G609G^ mice by improving WAT function, metabolic profile, and increasing body weight in ghrelin-treated HGPS mice. Our findings provide compelling *in vitro* and *in vivo* evidence that ghrelin ameliorates the characteristic changes caused by progerin-induced premature aging. Ghrelin ameliorates healthspan and extends the lifespan of these mice by improving white adipose tissue structure and function, rescuing the lipodystrophy phenotype. Given the multitude of actions attributed to ghrelin and the fact that ghrelin and its analogues have already been used in several clinical trials as a therapeutic strategy for the treatment of conditions such as of cachexia in chronic heart failure, cancer, end-stage-renal-disease or cystic fibrosis, frailty in the elderly, anorexia nervosa, growth hormone deficient patients, and sleep-wake regulation (80, 81), the findings of this study provide ghrelin as an effective intervention to delay premature aging in HGPS and normal cellular aging, thus enhancing healthspan and lifespan.

## Materials and Methods

### Cell cultures

Primary human dermal fibroblast cell cultures derived from two HGPS patients (HGADFN003, a 3-year-old male; and HGADFN127, a 2-year-old female) from the Progeria Research Foundation and were used as an *in vitro* model of HGPS. Primary cultures of human dermal fibroblasts obtained from a skin biopsy of a healthy individual (Control fibroblasts) were also used and provided by Coriell Cell Repositories. The cells were cultured in Dulbecco’s modified Eagle’s medium (DMEM, 4.5 g.L^-1^ D-glucose; Sigma) supplemented with 15 % fetal bovine serum (FBS; Gibco), 2 mM L-glutamine (Gibco) and 100 U/mL penicillin and 100 μg.mL^-1^ streptomycin (Gibco) and maintained at 37 ºC and 5 % CO2/air. All results shown in this article correspond to HGPS fibroblasts HGADFN003. We used in previously established non-HGPS control dermal fibroblast cultures in parallel. Control fibroblasts were used between passages 14 and 19, whereas HGPS fibroblasts were between 13 and 23.

### Experimental conditions

HGPS and Control fibroblasts were treated with human Ghrelin (1 nM; Bachem) every other day for up to one week, unless otherwise indicated. The lysosomal protein degradation inhibitor chloroquine (ChQ; 100 μM; Sigma-Aldrich) was added to the cell culture medium 30 minutes prior to ghrelin treatment for 6 hours.

### Autophagic flux measurement by LC3B turnover assay

Microtubule-associated protein light chain-3B (LC3B) is now the most widely used autophagy markers (82). Upon autophagy induction, the soluble cytosolic form of LC3B (LC3B-I) is conjugated to phosphatidylethanolamine to form a lipidated form (LC3B-II). LC3B-II becomes membrane-bound to the phagophores, autophagosomes and autolysosomes. LC3B-II is localized both in the luminal and cytosolic sites of the autophagic structures and undergoes degradation within the lysosome. Thus, LC3B-II’s steady-state levels and turnover within lysosomes can be used to measure autophagic activity in cells. However, because autophagy is a highly dynamic process, it should be noted that the amount of LCB3-II at a given time point does not necessarily estimate the autophagic flux. Therefore, it is important to measure the amount of LC3B-II delivered to lysosomes (82). Autophagic flux is often inferred based on LC3-II turnover, measured by Western blotting, in the presence and absence of lysosomal degradation. The relevant parameter in LC3B turnover assays is the difference in the amount of LC3B-II in the presence and absence of saturating levels of inhibitors, which can be used to examine the transit of LC3B-II through the autophagic pathway and its accumulation within lysosomes. This assay will reflect the net amount of LC3B-II delivered to lysosomes, which will be the autophagy flux measure. If the flux occurs, the amount of LC3B-II can be prevented using compounds that neutralize the lysosomal pH, such as chloroquine or NH4Cl, inhibiting lysosomal degradation (82). In our study, we determined autophagic flux by measuring the amount of LC3B-II delivered to the lysosomes by comparing the LC3B-II amount in the presence and absence of the lysosomal inhibitor chloroquine (ChQ; 100 μM) by Western blotting. Chloroquine was added to the cell culture medium 30 minutes before the addition of ghrelin (6 hours-treatment). For each experimental condition, untreated HGPS cells (used as control) and ghrelin-treated HGPS cells, autophagic flux, expressed as “LC3B-II net flux”, was determined by subtracting the densitometric value of the LC3B-II band of the chloroquine-untreated sample (ChQ-LC3B-II) from the densitometric value of the LC3B-II band of the corresponding chloroquine-treated sample (ChQ+LC3B-II). For each independent experiment, the values obtained upon subtraction for each condition were normalized to the control condition (untreated HGPS cells). The results are represented as mean values for each experimental condition. The LC3B-net flux results are shown in parallel with the results regarding the quantification of the steady-state levels of LC3B-II (densitometric values of LC3B-II are normalized to the loading control β-tubulin) in the presence or absence of chloroquine in untreated (HGPS cells) and ghrelin-treated cells to assess the effect of ghrelin on LC3B-II levels as well as demonstrate the accumulation of LC3B-II levels upon chloroquine treatment.

### Immunocytochemistry

Immunocytochemistry was performed as described in the *SI Appendix, Material and Methods*.

### Senescence-associated-β-galactosidase (SA-β-Gal) assay

To measure senescence-associated-β-galactosidase (SA-β-Gal) activity in HGPS cells, the medium was removed, and cells were washed twice with PBS at room temperature. The cells were then fixed with ice-cold 4 % paraformaldehyde for 3 minutes, followed by two rinses with PBS at room temperature. Next, cells were exposed to fresh SA-β-Gal staining solution containing 1x citric acid/sodium phosphate buffer (pH 6.0), 5 mM potassium ferrocyanide, 5 mM potassium ferricyanide, 150 mM NaCl, 2 mM MgCl2, and X-Gal (1 mg.mL^-1^; Invitrogen Molecular Probes) in water. The cells were incubated overnight in a non-CO2 incubator, at 37 ºC. Afterward, the staining solution was washed away, and cells were rinsed twice with PBS. Finally, the coverslips were mounted on glass slides with an Aqua-Polymount mounting medium (Polysciences, Inc.). To quantify SA-β-Gal-positive cells, 30 randomly chosen non-overlapping fields were examined for each coverslip of each experimental condition at x200 magnification. The number of positive cells was directly counted.

### Mouse strains and experiments

The mouse model of HGPS carrying the *LMNA* causative mutation p.Gly609Gly (*Lmna*^G609G/G609G^, with the genetic background of C57BL/6) were used to perform *in vivo* experimental studies. The infertile *Lmna*^G609G/G609G^ mice were obtained by crossing heterozygous (*Lmna*^G609G/+^) females and males. Both male and female animals were used in all experiments. Animals were housed in pairs or fours *per* cage, under a 12-hour:12-hour light/dark cycle, with controlled temperature and humidity and *ad libitum* access to normal chow diet. All experimental work was approved by CNC Animal Welfare Body (ORBEA 329 and DGAV 009428) and performed following the European Community directive for the care and use of laboratory animals (86/609/EEC) and the Portuguese law for the care and use of experimental animals (Decree-law 113-2013). The animals were housed in the licensed animal facility of CNC International Animal Welfare Assurance number 520.000.000.2006). Animal experimentation was performed by credited and trained investigators, as required by the Portuguese authorities. The study was included in projects approved and financed by the Progeria Research Foundation, which approved the use of animals for this study (PRF2014-53 and PRF2015-60). Ghrelin (Bachem) was administered at a concentration of 50 μg/Kg each in 0.9 % NaCl through daily subcutaneous injection for 6-weeks. Control mice were administered the same routine as a vehicle, 0.9 % NaCl. Survival curve analysis, performed twice (*N*=9-13), was terminated when animals reached predefined humane endpoints according to the FMUC/CNC-UC animal facility.

### Food intake and body weight analysis

All experiments measured body weight and food intake twice a week. Body weight gain was calculated as a percentage of weight gain and presented in a plot graph as Body Weight (% of Initial Body Weight). Food intake was presented as the total ingestion of calories within the experimental period. Since mice were not kept in individual cages due to rules respecting our animal facility, the individual food intake was calculated as follows: (Total food intake per cage/ Total weight *per* cage) X Mouse weight (g) (83).

### Tissue and blood collection

Animals were euthanized 6-weeks after starting treatment by sodium pentobarbital overdose. Animals from each experimental group were randomly distributed for blood collection, and peripheral organs and tissues were collected for RNA and/or protein extraction and/or histological analysis. Blood was collected upon decapitation, and serum was obtained by centrifugation (2,000×g for 15 minutes). After decapitation, several organs, and tissues, such as gonadal white adipose tissue (WAT) and interscapular brown adipose tissue (BAT), skin, heart, aorta, liver, skeletal muscle, and spleen, were collected and weighed. These organs were then cut and divided. A part of each organ was kept at -80 °C for protein and RNA extraction purposes, while the other was kept in a 10 % neutral buffered formalin solution for 48 hours to prepare them for histological processing. Serum samples were kept at -20 °C until use.

### Biochemical parameters assessment

Biochemical profiles were measured on Cobas 6000 from Roche (Basel, Switzerland). Glucose levels were measured using a FreeStyle Precision Neo glucometer (Abbott, Chicago, IL, USA).

### Plasma metabolism multiplex assay array

Metabolic hormone levels were measured in mouse serum samples replicates using a MILLIPLEX® Mouse Metabolic Hormone Magnetic Bead Panel, Metabolism Multiplex Assay Array (#MMHMAG-44K; Millipore). The levels were determined through a multiplex array using the Bio-PlexTM system (a service provided at 476 Biocant Park, Cantanhede, Portugal).

### Gene expression analysis

Total RNA extraction, reverse transcription, and quantitative real-time PCR (qRT-PCR) analysis were performed as described in the *SI Appendix, Material and Methods*.

### Western blotting

Western blotting was performed as described in *SI Appendix, Material and Methods*.

### Histological analysis

Tissue samples were collected and fixed in 10 % formalin, cut into small fragments, and underwent several steps for paraffin embedding: ethanol 70 % for 1 hour; two series of ethanol 95 %, 40 minutes each; two series of ethanol 100 %, 1 hour each; two series of xylene, 1 hour each and two series of paraffin at 56 ºC, 1 hour each. At the end of this process, tissue samples were included in paraffin blocks. Paraffin blocks were sectioned using a microtome (HM325, Thermo Fisher Scientific), and 3-5 μm thickness sections were placed into microscopy slides until use. Haematoxylin-eosin staining was performed according to standard procedures. After staining, the sections were mounted on slides with Richard-Allan Scientific Mounting Medium (Thermo Fisher Scientific). The nuclei were stained blue, and the cytoplasm red to detect structural alterations in the tissue. Masson’s trichrome (Thermo Fisher Scientific) and Picro-Sirius Red (Direct Red 80 dye/Sirius Red F3BA; Sigma Aldrich) staining were performed according to the manufacturer’s protocol. Tissue section images were acquired using a Zeiss Axio Imager Z2 microscope (Zeiss) with the Plan-Apochromat 20x/0.8 M27 objective. The images were analyzed with FIJI (Fiji is Just ImageJ) Software. The analysis was performed by a researcher who was unaware of the experimental groups.

### Immunohistochemistry

Immunohistochemistry was performed according to the methods described in the *SI Appendix, Material and Methods*. Protein levels of Progerin, alpha-smooth muscle actin (α-SMA), Ki-67, and keratin 1 (KRT1) were accessed by fluorescent immunohistochemistry of formalin-fixed, paraffin-embedded skin and/or aorta sections collected on Superfrost slides. Similarly, to the histological staining protocol described above, an initial step of deparaffinization and rehydration was performed by successively immersing skin sections in xylene (2×5 minutes), 100 % (v/v) ethanol (2×5 minutes), 95 %, 70 % and 50 % (v/v) ethanol (1×3 minutes each), followed by a 5-minute wash in PBS. An antigen retrieval step was performed to expose the antigenic sites that may have been masked by formalin fixation. The skin and/or aorta sections were incubated with sodium citrate buffer (pH 6.0), heated in a water bath at 100 °C for 20 minutes, and allowed to cool down for 1 hour. Sections were then washed with PBS (3×2 minutes). To block nonspecific antibody binding sites and permeabilize the sample, sections were outlined with a hydrophobic pen and incubated with a blocking solution 3 % BSA/10 % goat serum/ 0.1 % triton X-100/ PBS for 1 hour at room temperature. Primary antibodies used were mouse anti-Progerin (1:250; Sigma), rabbit anti-Ki-67 (1:250; Abcam), rabbit anti-KRT1 (1:1,000; BioLegend) and mouse anti-alpha-smooth muscle actin Cy3 (1:250; Sigma), diluted in the blocking solution, overnight at 4 °C. Antibody–antigen complexes were detected using secondary antibodies Alexa Fluor 568 conjugated goat anti-rabbit or goat anti-mouse IgG for 2 hours at room temperature. Nuclei counterstaining was performed with Hoechst 33342 (2 μg.mL^-1^; Invitrogen Molecular Probes) during secondary antibody incubation. Lastly, sections were rinsed three times with PBS and the coverslips were mounted on glass slides with Mowiol® 4-88 (Sigma) mounting medium. Images were acquired using an Axio Observer Z1 microsocpe (Carl Zeiss) using a Plan-Apochromat 20x/0.8 M27 objective. Images were analysed with FIJI (Fiji is Just ImageJ) Software or QuPath Software. The analysis was performed by a researcher who was unaware of the experimental groups.

### Statistical analysis

The results are expressed as mean±SEM. Statistical analyses were conducted using one-way ANOVA followed by Tukey’s multiple comparison test or unpaired Student’s *t*-test, depending on the number of experimental groups in each experimental condition. Prism 8.4.2 (GraphPad Software) was used for the statistical analysis. The statistical parameters can be found in the figure legends.

## Acknowledgments

This work was financed by: Progeria Research Foundation (PRF2014-53 and PRF2015-60), the European Regional Development Fund (ERDF), through the Centro 2020 Regional Operational Programme projects CENTRO-01-0145-FEDER-000012 (HealthyAging 2020), CENTRO-01-0246-FEDER-000010 (MIA-Portugal), and COMPETE 2020 – Operational Programme for Competitiveness and Internationalisation; European Union’s Horizon 2020 research and innovation programme (grant agreement N .º 857524); FCT – Fundação para a Ciência e a Tecnologia, under project[s] POCI-01-0145-FEDER-030167 (NiNjA), UIDB/04539/2020, LA/P/0058/2020, FCT Investigator Programme (IF/00825/2015), SFRH/BD/120023/2016, COVID/BD/152130/2021.. C.L-O. is supported by grants from the European Research Council (ERC Advanced Grant, DeAge), and Ministerio de Ciencia e Innovación, Spain.

