## Supplemental Information for "Ghrelin delays premature aging in Hutchinson-Gilford progeria syndrome"

[1]CNC - Center for Neuroscience and Cell Biology, University of Coimbra, 3004-504 Coimbra, Portugal
[2]CIBB - Center for Innovative Biomedicine and Biotechnology, University of Coimbra, 3004-504 Coimbra, Portugal
[3]Departamento de Bioquímica y Biología Molecular, Facultad de Medicina, Instituto Universitario de Oncología, Universidad de Oviedo, 33006 Oviedo, Spain
[4]CECS, I-Stem, 91100 Corbeil-Essonnes, France
[5]INSERM U861, I-Stem, 91100 Corbeil-Essonnes, France
[6]UEVE U861, I-Stem, 91100 Corbeil-Essonnes, France
[7]Faculty of Pharmacy, University of Coimbra, 3000-548 Coimbra, Portugal
[8]MIA-Portugal - Multidisciplinar Institute of Ageing, University of Coimbra, 3000-370 Coimbra, Portugal
[#]Equal senior contribution

*Célia A. Aveleira & Cláudia Cavadas

**Supplementary Figure S1 Progerin mRNA expression was not altered by ghrelin in HGPS fibroblasts.** (*A* and *B*) HGPS fibroblasts were exposed to ghrelin (1 nM) for 6 hours (HGPS+Ghrelin) (*A*), or for one week, treated every other day (*B*). HGPS-untreated cells were used as control (HGPS). Quantitative polymerase chain reaction analysis of progerin mRNA levels in HGPS fibroblasts upon 6 hours (*N*=4) or 1 week (*N*=5) of ghrelin treatment. Data are expressed as the mean±SEM of, at least, three independent experiments, and are expressed as a percentage of HGPS. HGPS = Hutchinson-Gilford progeria syndrome.

### Figure S1

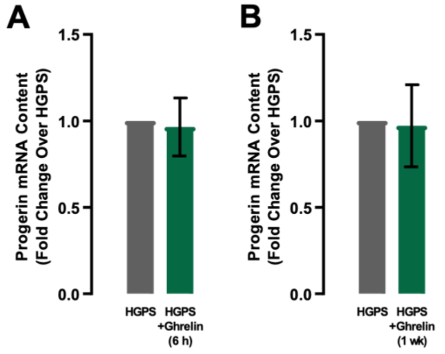

**Supplementary Figure S2 Ghrelin rescues nuclear morphology and increases cell proliferation in control fibroblasts.** (*A*-*E*) Control fibroblasts were exposed to ghrelin (1 nM; Control+Ghrelin) for one week. Untreated cells were used as control (Control). (*A*) Control fibroblasts were immunolabeled for Lamin A/C (red) and nuclei were stained with Hoechst 33342 (blue, bottom panel). Images are representative of four independent experiments. Scale bar, 10 µm. Quantification of the number of misshapen nuclei (*B*) and nuclear circularity (*C*) upon ghrelin treatment. An equal number of nuclei (>400) were randomly analyzed for each condition. Circularity (defined as 4*π*area/perimeter^2) was measured using ImageJ. A circularity value equal to 1 corresponds to perfectly circular nuclei. (*D*) Ghrelin increases cell proliferation, as determined by Ki-67 immunoreactivity. Cells were immunolabeled for Ki-67 (red) and nuclei were stained with Hoechst (blue). Representative images of five independent experiments are shown. Scale bar, 10 µm. (*E*) Quantification of the number of Ki-67-positive cells in untreated and ghrelin-treated Control cells. The results represent the mean±SEM of five independent experiments and are expressed as a percentage of Control. **P*<0.05 and ***P*<0.01, significantly different from Control, as determined by Student's *t*-test.

### Figure S2

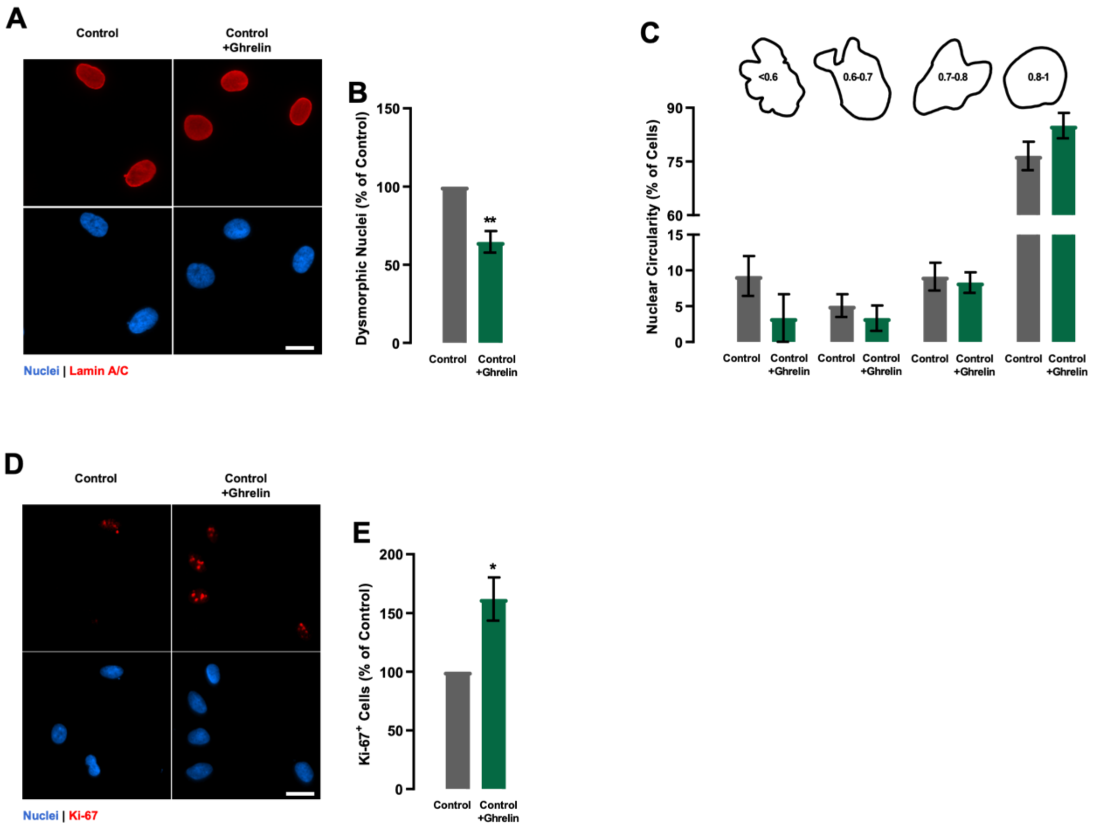

**Supplementary Figure S3 Amelioration of age-related alterations in the periphery by ghrelin treatment in *Lmna*$^{G609G/G609G}$ mice.** (*A*) Organ weight expressed as a percentage of % of body weight in 3 months-old vehicle- and ghrelin-treated *Lmna*$^{+/+}$ and *Lmna*$^{G609G/G609G}$ mice. (*B*) Representative images of Hematoxylin-eosin- (top panel) and Masson's Trichrome (bottom panel) stained sections of liver of vehicle- and ghrelin-treated *Lmna*$^{+/+}$ and *Lmna*$^{G609G/G609G}$ mice. Scale bar, 100 µm. (*C*) Representative images of Hematoxylin-eosin-stained spleen sections of vehicle- and ghrelin-treated *Lmna*$^{+/+}$ and *Lmna*$^{G609G/G609G}$ mice. Scale bar, 100 µm. (*D*) Quantification of spleen white pulp area, expressed as % of total area, in the spleen, respectively, of vehicle- and ghrelin-treated *Lmna*$^{+/+}$ and *Lmna*$^{G609G/G609G}$ mice. (*E*) Representative images of Hematoxylin-eosin-stained skeletal muscle sections of vehicle- and ghrelin-treated *Lmna*$^{+/+}$ and *Lmna*$^{G609G/G609G}$ mice. Scale bar, 100 µm. (*F*) Quantification of muscle fiber cross-sectional area expressed in µm$^2$. Data are expressed as the mean±SEM. *N*=5-12 *per* group. *$P<0.05$ and ****$P<0.0001$, significantly different from *Lmna*$^{+/+}$ mice; $^{\#}P<0.05$ and $^{\#\#}P<0.01$, significantly different compared to *Lmna*$^{G609G/G609G}$ mice, as determined by analysis of variance, followed Tukey's multiple comparison test.

### Figure S3

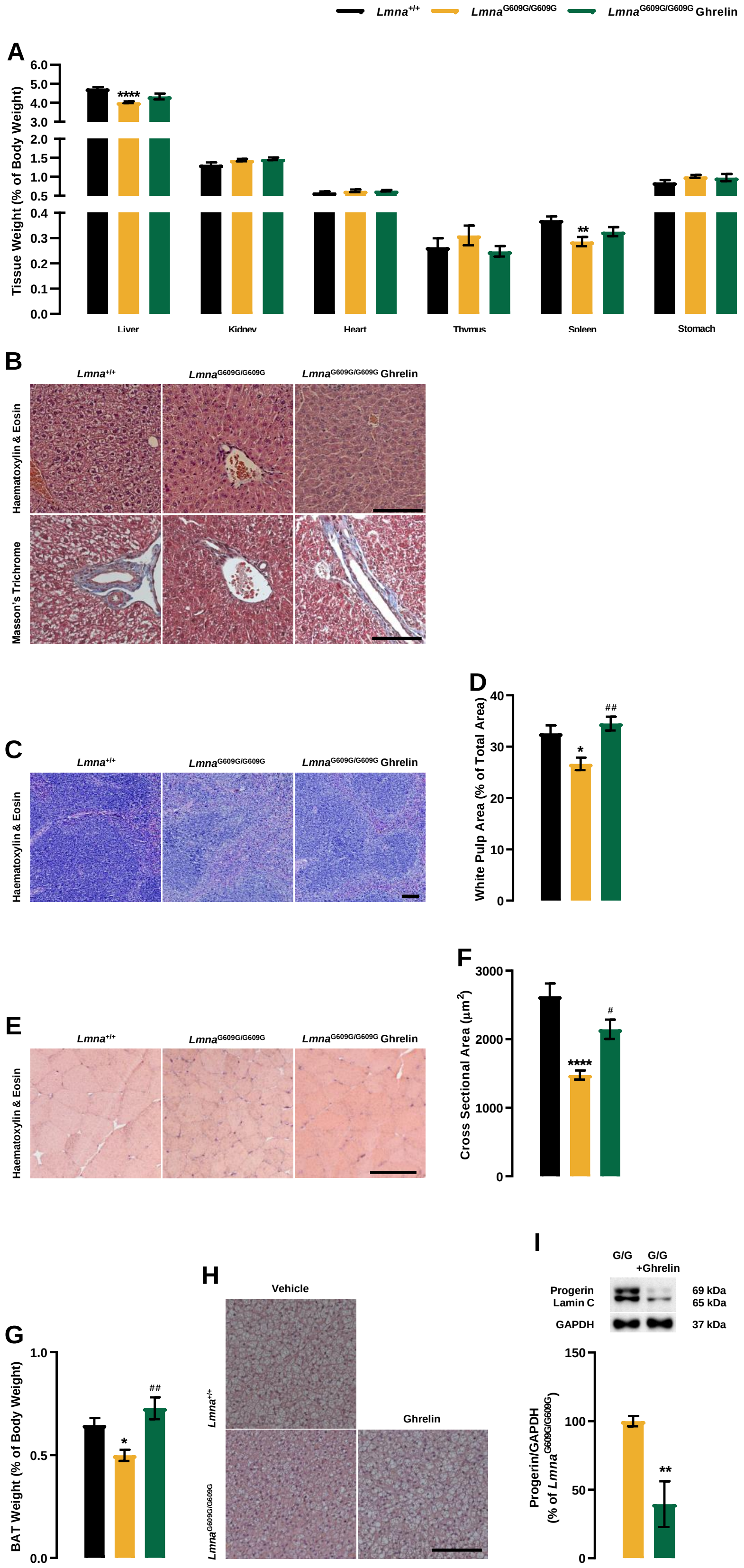

Lmna+/+
LmnaG609G/G609G
LmnaG609G/G609G Ghrelin
A
Tissue Weight (% of Body Weight)
6.0
5.0
4.0
3.0
2.0
1.5
1.0
0.5
0.4
0.3
0.2
0.1
0.0
****
**
Liver
Kidney
Heart
Thymus
Spleen
Stomach
B
Haematoxylin & Eosin
Masson's Trichrome
C
D
White Pulp Area (% of Total Area)
40
30
20
10
0
*
##
E
F
Cross Sectional Area (μm2)
3000
2000
1000
0
****
#
G
BAT Weight (% of Body Weight)
1.0
0.5
0.0
*
##
H
Vehicle
Ghrelin
I
G/G
G/G +Ghrelin
Progerin
Lamin C
GAPDH
69 kDa
65 kDa
37 kDa
Progerin/GAPDH (% of LmnaG609G/G609G)
150
100
50
0
**

**Supplementary Table S1. Summary of primers' sequences used for gene expression analysis and optimal conditions for qRT-PCR.**

| *GENE* | Primer Sequence (5' → 3') | *Function* | Ta (ºC) |
|---|---|---|---|
| *PROGERIN* | F:CTCAGGAGCCCAGAGCC<br>R:GGCATGAGGTGAGGAGGAC | HGPS mutant protein | 59 ºC |
| *ACTB* | F:TGATCTTGATCTTCATTGTG<br>R:AACTACCTTCAACTCCATC | Housekeeping gene | 59 ºC |
| *HPRT* | F:GGCTTATATCCAACACTTCG<br>R:TGACACTGGCAAAACAATG | Housekeeping gene | 59 ºC |
| *Progerin* | F:CGCTGAGTACAACCTGCG<br>R:TGGCAGGTCCCAGATTACAT | HGPS mutant protein | 58 ºC |
| *Lep* | F:TTTCACACACGCAGTCGGTA<br>R:GGACCTGTTGATAGACTGCCA | Adipose tissue function | 60 ºC |
| *Adipoq* | F:TGTCCCCATGAGTACCAGACT<br>R:TCCTGAGCCCTTTTGGTGTC | Adipose tissue function | 58 ºC |
| *Slc2a4* | F:CCGGACCCTATACCCTATTCA<br>R:GGGTTCCCCATCGTCAGA | Adipose tissue function | 60 ºC |
| *Cebpb* | F:AATCCGGATCAAACGTGGCT<br>R: CCGCAGGAACATCTTTAAGT | Adipogenic early differentiation regulator | 57ºC |
| *Cebpd* | F:CGGCCTTCTACGAGCCAG<br>R:GTCGTACATGGCAGGAGTCG | Adipogenic early differentiation regulator | 59 ºC |
| *Pparg* | F:GCCTATGAGCACTTCACAAGAAAT<br>R:AGTGGTCTTCCATCACGGAG | Adipogenic late differentiation regulator | 58 ºC |
| *Cebpa* | F:TACCGAGTAGGGGGAGCAAA<br>R:TCATTTTTCTCACGGGGCCA | Adipogenic late differentiation regulator | 60 ºC |
| *Pepck* | F:CCAACGTGGCCGAGACTAGCG<br>R:GGCACATGGTTCCGCGTCCT | Gluconeogenesis | 60 ºC |
| *Srebp-1c* | F:GATCAAAGAGGAGCCAGTGC<br>R:TAGATGGTGGCTGCTGAGTG | Cholesterol and fatty acid biosynthesis | 58 ºC |
| *Acc* | F:GGAGATGTACGCTGACCGAGAA<br>R:ACCCGACGCATGGTTTTCA | Fatty acid biosynthesis | 60 ºC |
| *FasN* | F:AGCTTCGGCTGCTGTTGGAAGT<br>R:TCGGATGCCTCTGAACCACTCACA | Fatty acid biosynthesis | 60 ºC |
| *Scd1* | F:CCGGAGACCCCTTAGATCGA<br>R:TAGCCTGTAAAAGATTTCTGCAAACC | Fatty acid oxidation | 60 ºC |
| *Cpt1* | F:GGTTGCTGATGACGGCTATGGTGT<br>R:GCGGTGAGGCCAAACAAGGTGATA | Fatty acid oxidation | 60 ºC |
| *Acly* | F:GCCAGCGGGAGCACATC<br>R:CTTTGCAGGTGCCACTTCATC | Fatty acid biosynthesis | 60 ºC |
| *Mcad* | F:AACACTTACTATGCCTCGATTGCA<br>R:CCATAGCCTCCGAAAATCTGAA | Fatty acid oxidation | 60 ºC |
| *Hprt* | F:GCTTACCTCACTGCTTTCCG<br>R:CATCATCGCTAATCACGACGC | Housekeeping gene | 58 ºC |

***F:*** Primer forward sequence; ***R:*** Primer reverse sequence; ***Ta (ºC):*** Annealing temperature

**Supporting Material and Methods**

**Immunocytochemistry.** After a 1-week treatment with ghrelin, cells were washed twice with PBS (pH 7.4) at 37 ºC and then fixed in ice-cold 4 % paraformaldehyde for 15 minutes. Cells were then rinsed three times with ice-cold PBS and permeabilized with 0.1 % (v/v) TX-100/PBS for 10 minutes at room temperature. Cells were then washed twice with PBS and blocked with 3 % BSA/10 % goat serum/ PBS for 1 hour at room temperature. Afterwards, cells were incubated overnight at 4 ºC with primary antibodies. The primary antibodies used were mouse anti-Progerin (1:500; Sigma), mouse anti-Lamin A/C (1:500; Millipore), mouse anti-γ-H2AX (1:500; Millipore) and mouse anti-Ki-67 (1:400; Abcam). After incubation, cells were washed three times for 5 minutes with PBS and incubated with the respective secondary antibody or 1 hour at room temperature. The secondary antibodies used were Alexa-Fluor 488- or Alexa Fluor 568 conjugated goat anti-mouse IgG. The nuclei were stained with Hoechst 33342 (2 μg.$mL^{-1}$; Invitrogen Molecular Probes) during secondary antibody incubation. Lastly, cells were rinsed three times with PBS and the coverslips were mounted on glass slides with Aqua-Polymount (Polysciences, Inc.) mounting medium. Quantification of Ki-67-positive cells and misshapen/blebbing nuclei was done by direct counting of positive cells in 40 randomly chosen, non-overlapping fields (x400 magnification), normalized to the total number of nuclei stained with Hoechst 33342 for each coverslip of each experimental condition. Results represent the mean±SEM of four independents experiments (at least 400 cells analyzed for each experimental condition) and are expressed as percentage of non-treated cells. The analysis of nuclei morphology parameters was performed using FIJI (Fiji is Just ImageJ) Software, using a homemade macro. Briefly, images were thresholded and nuclei were automatically detected based on their size (100-infinite) and circularity (0.3-1). Then, all images were manually reanalyzed to confirm the automatic selection. Finally, the following parameters were measured for each selected nucleus: area, perimeter, and circularity (4π*area/perimeter^2). For γ-H2AX *foci* quantification 20 randomly chosen non-overlapping z-stacking image were used for each experimental condition. First, the nuclei in all images were automatically detected using the nuclear morphology parameters macro above described. Then, the FindFoci plugin was applied in order to identify the peak corresponding to each *foci* in each nuclei, setting the minimum peak size above saddle on 5 (1). The number of *foci*/nuclei were calculated for each nucleus and condition.

**Gene expression analysis**. Total RNA was extracted from HGPS cells using the RNeasy Mini Kit (Qiagen) according to the manufacturer's instructions. Briefly, cells were lysed, the total RNA was adsorbed to a silica matrix, washed with the recommended buffers, and eluted with 30 µL of RNase-free water by centrifugation. RNA samples were treated with RNase-free DNAse (Qiagen) to exclude any contamination with genomic DNA. Total RNA from WAT tissue was isolated from snap-frozen samples, then lysed and homogenized with TRI Reagent (Sigma-Aldrich; 1 mL of TRI Reagent per 100 mg of tissue) using the pestle. To ensure complete dissociation of tissue, all samples were left at room temperature for 5 minutes. Subsequently, samples were centrifuged for 10 minutes at 12,000 x g at 4 ºC for the deposition of the insoluble material. The supernatant was collected to a respective new tube and 0.2 mL of chloroform (Sigma-Aldrich) was added. The samples were then shaken vigorously and left at room temperature for 3 minutes. Samples were once again centrifuged at 12,000 x g for 15 minutes at 4 ºC, and the mixture was separated into 3 phases: a red organic phase, which contains the protein, an interphase, which contains DNA, and a colorless upper aqueous phase, which contains RNA. After isolating the aqueous phase, RNA was purified using the NucleoSpin RNA Kit (Macherey-Nagel) following the instructions of the manufacturer. Briefly, RNA-containing aqueous fractions were mixed with an equal volume of 70 % (v/v) ethanol and then loaded on a NucleoSpin® RNA Column for RNA binding to the silica membranes. RNA was washed with the supplied buffers and eluted in 40 µL of RNase free water by centrifugation. RNA samples were treated with RNase free DNase (Macherey-Nagel) to avoid contamination with genomic DNA. The total amount of RNA was quantified by optical density measurements (OD) using a ND-1000 nanodrop spectrophotometer (Thermo Scientific), and purity was assessed by measuring the ratio of OD at 260 and 280 nm.RNA samples were stored at -80 °C until use. Reverse transcription into cDNA was performed using the iScript cDNA Synthesis Kit (Bio-Rad) according to the manufacturer's instructions. Briefly, 1 µg total RNA from each sample was transcribed into cDNA in a 30 µL reaction with 1x iScript reaction buffer and 1 µL iScript reverse transcriptase. Reverse transcription reactions were performed in a thermal cycler at 25 °C for 5 minutes, 46 °C for 30 minutes, 95 °C for 5 minutes, and 4 °C for 5 minutes. cDNA samples were then stored at -20 °C until use. mRNA expression was measured by qRT-PCR in the StepOnePLusTM Real-Time PCR System (Applied Biosystems) using 96-well optical plates (Thermo) and SsoAdvancedTM Universal SYBR® Green Supermix (BioRad). A master mix was prepared for each primer set, containing the appropriate volume of 2× SsoAdvancedTM Universal SYBR® Green Supermix and 500 nM of each specific gene primer. For each reaction, 6 µL of the master mix was added to 4 µL of template cDNA. All reactions were performed in duplicates (two cDNA reactions *per* RNA sample). Negative controls, including no template control (NTC) and no reverse transcriptase control (NoRT) were used. Reactions were performed according to the manufacturer's recommendations: 95 °C for 3 minutes, followed by 40 cycles at 95 °C for 5 seconds and 59 °C for 15 seconds. The melting curve protocol started immediately after amplification. The StepOnePLus Software (Applied Biosystems) automatically determined the amplification efficiency for each gene and threshold cycle (Ct) determination. Relative mRNA quantification was performed using the ΔCt method for genes with the same amplification efficiency. The sequences of the primers used are listed in *SI Appendix*, Table S1.

**Western blotting.** Protein samples were collected from cell pellets resuspended in (RIPA) buffer (50 mM Tris-HCl, pH 7.4; 150 mM NaCl; 5 mM EDTA; 1 % Triton X-100; 0.5 % deoxycholate; 0.1 % sodium dodecyl sulphate (SDS); 200 µM phenylmethylsulphonylfluoride (PMSF); 1 mM dithiothreitol (DDT); 1 mM $Na_3VO_4$; 10 mM NaF) supplemented with Complete™ Mini Protease Inhibitor. Extraction of tissue protein was performed after isolating the organic phase. Briefly, 100 % ethanol was added to precipitate DNA. Tubes were mixed by inversion and centrifuged at 2,000xg for 10 minutes, at 4 °C. The phenol-ethanol supernatant was removed to 2 mL tubes, for protein extraction. Protein was precipitated with isopropanol, followed by mixing and 10 minutes incubation at room temperature. Subsequently, samples were centrifuged at 12,000xg for 10 minutes and the supernatant discarded. Protein pellets were next washed twice with 0.3 M guanidine hydrochloride in 95 % ethanol. In each wash, tubes were vigorously shaken, incubated at room temperature for 20 minutes followed by centrifugation at 7,500xg for 5 minutes at 4 °C. After the final wash and spin, 100 % ethanol was added, and samples were incubated at room temperature for 20 minutes, followed by a final centrifugation at 7,500xg for 5 minutes at 4 °C. The supernatant was removed and 10 M Urea/ 500 nM DTT was added to the protein pellets. Protein content was determined using the bicinchoninic acid (BCA) protein assay (Thermo Scientific), following to the manufacturer’s instructions. After protein quantification, the samples were then boiled in SDS-sample-buffer, and equal amounts of protein extracts were run on 4–10 % or -12 % polyacrylamide gel (Bio-Rad) and transferred onto an Immobilon-P polyvinylidene fluoride (PVDF) membrane (Merck Millipore). The membrane was blocked by 5 % BSA or milk in TBST (TBS with 0.1 % Tween-20) at room temperature for 1 hour and then incubated with primary antibodies in 1x TBST with 5 % BSA or milk overnight. The primary antibodies used were mouse anti-laminA/C (1:1000; Developmental Studies Hybridoma Bank; MANLAC1 (4A7)), mouse anti-p53 (1:500; Abcam), rabbit anti-p21 (1:500; Santa Cruz Biotechnology), and rabbit anti-LC3B, anti-MTOR, and anti-phospho-MTOR (Ser2448) (all at a dilution of 1:1,000; from Cell Signaling). The secondary antibodies used were rabbit or mouse IgG-specific alkaline phosphatase-linked (all from PIERCE), in a dilution of 1:10,000 in the same blocking solution as the respective primary antibody, and signal was detected by enhanced chemifluorescence (ECF, GE Healthcare, Little Chalfont, UK) in a VersaDoc Imaging System (Bio-Rad, California, USA). The optical density of the bands was quantified using Quantity One Software (Bio-Rad, California, USA). The results were normalized to the amount of β-Tubulin or β-Actin and or GAPDH (all at a dilution of 10,000) and are expressed as the relative amount compared with control.
